# Deciphering lung microbial EV pneumotypes in acute respiratory distress syndrome proteome wide

**DOI:** 10.64898/2026.11.24.767187

**Authors:** Ningshan Li, Runzhen Zhao, Arslan Bisharat, David Smith, Emily Gilbert, Jiwang Zhang, Hong-Long Ji

## Abstract

**Background:** Pneumotypes hold potential predictive value for pathogenesis, outcomes, and interventions. Lung microbiome releases extracellular vesicles (EVs). However, EV-derived pneumotypes in acute respiratory distress syndrome (ARDS) have not been studied yet.

**Objectives:** We aimed to identify pneumotypes in ARDS patients by comprehensively profiling lung microbiome-released EV proteins in bronchioalveolar lavage fluid (BALF).

**Methods:** We extracted EVs from 106 participants (57 cases and 49 healthy controls) enrolled in the ARDSNet and SPIROMICS clinical trials and subjected to nanoparticle liquid chromatography-mass spectrometry (LC-MS) to create microbial metaproteomics. Differentially expressed proteins (DEPs) were identified by Wilcoxon rank-sum test, and spectral clustering of these DEPs was utilized to reveal patterns in microbial EV protein profiles. The association between pneumotypes and the disease was assessed. Pneumotype-associated DEPs were identified using logistic regression, and their functional annotation was performed based on the eggNOG 5 database. The taxonomy of lung microbes was determined at the species-level and ranked by biomass contributions.

**Results:** Ninety DEPs were identified in ARDS lavage samples (53 upregulated and 37 downregulated) compared with healthy controls. There were five pneumotypes, namely, P1, P2, P3, P4, and P5, identified by clustering the differentially expressed microbial EV proteins. Few pneumotype-associated DEPs were shared among these pneumotypes and mainly correlated with ARDS etiology and organ functions. The significant difference in mortality was observed in P3 and P4 subtypes. In contrast, all patients in P1 and P5 subgroups survived. Patients in P2 and P3 groups experienced a significant reduction in both of central nervous system (CNS) failure-free days and the days without organ failure compared with other subgroups. Besides, patients in P3 and P4 exhibited a higher prevalence of baseline shock and hypertension, greater use of vasopressors or inotropes within the previous 24 hours, higher positive end-expiratory pressure during ventilation, and lower body temperature and hematocrit levels. Alpha and beta diversity analyses showed significant shifts in P2 patients. The taxonomy of the top-ranked microbes exhibited different pathogens in P2, P3, and P4.

**Conclusions:** This study, for the first time, identified five EV expression patterns of lung microbiota in the BALF of ARDS patients. These intrinsically featured EV pneumotypes in the lung are associated with mortality, etiology, lung functions, medication administration, and disease severity in ARDS patients.

## 1. Background

**Acute respiratory distress syndrome** (ARDS) is a heterogeneous critical condition with high mortality and frequently occurs in intensive care unit (ICU) patients. The leading etiology of ARDS is pneumonia predominately caused by local and systemic infections. The high mortality rate has been attributable to non-specific effective therapy, which is challenged by the heterogenicity of the disease. The lower respiratory tract microbiota is similar in healthy subjects to the oropharyngeal microbiota and dominated by members of the Firmicutes, Bacteroidetes, and Proteobacteria phyla. Lung microbiota can be changed by infectious and noninfectious respiratory diseases as well as life habits. Understanding the expression patterns of lung microbiota in respiratory diseases holds the potential to study microbe-host interactions, develop target therapies, and improve precision medicine (Sumner et al., 2025).

The study of pneumotypes in respiratory diseases is still in its infancy stage. The expression patterns of lung microbiota have been investigated in several pulmonary diseases, including noninfectious lung cancer, asthma, and chronic obstructive pulmonary disease (COPD) as well as infectious nontuberculous mycobacterial pulmonary disease and COVID-19 in adults (Roshni et al., 2023; Chen et al., 2025b; Cheng and Wang, 2025; Lin et al., 2025; Qiu and Li, 2025) and newborns (Morin et al., 2020; Brewer et al., 2021; Chu et al., 2021). In general, metagenomic sequencing approach was applied to characterize the lung microbiota in bronchoalveolar lavage fluid (BALF) and cells. Among these studies, reduced diversity and increased dysbiosis were associated with disease severity and other outcomes (Faner et al., 2017). However, the specific microbial determinants driving clinical outcomes in ARDS have not yet been studied.

This study aimed to identify pneumotypes in ARDS patients by clustering unbiased high throughput microbial extracellular vesicles (EVs) metaproteomic datasets. To the best of our knowledge, this proteome wide identification of five distinct expression patterns in the BALF biopsies is the first study of ARDS pneumotypes. These newly identified EV pneumotypes are characterized by diverse biological processes and clinical features, which pave a path to identify pathogenic mechanisms and improve personalized interventions in ARDS.

## 2. Materials & Methods

### 2.1 Study design

Our cohort comprised 57 ARDS patients and 49 healthy controls who participated in the ARDSNet (National Heart, 2011) and SPIROMICS clinical trials (Couper et al., 2014). Clinical variables and BALF samples collected by the trials were provided by the Biologic Specimen and Data Repository Coordinating Center (BioLINCC). Detailed descriptions of clinical variable cleaning, extracellular vesicle extraction, liquid chromatography-mass spectrometry (LC-MS)-based metaproteomic data generation, and preprocessing are provided in the Supplementary Methods. This study was reviewed and approved by the Institutional Review Board of Loyola University Chicago.

### 2.2 Differential analysis

Differential analysis was performed on the preprocessed metaproteomic data to identify differentially expressed proteins (DEPs) between ARDS cases and controls using the Wilcoxon rank-sum test (Wilcoxon, 1945). Proteins with a fold change greater than 4 and a false discovery rate (FDR) < 0.05 were considered as DEPs which were determined by the Benjamini-Hochberg (BH) procedure (Benjamini and Hochberg, 1995). The fold change threshold of 4 was selected to achieve greater than 80% statistical power under a two-sample Wilcoxon rank-sum test framework, as determined by *post hoc* power analysis using G*Power (version 3.1.9.7) (Faul et al., 2007; Faul et al., 2009), with a significance level of α = 0.05 and Cohen’s *d* (Cohen, 1988) effect size estimated from the median of the observed pooled standard deviations of log_2_-transformed protein intensities.

### 2.3 Clustering

Reporting of clustering follows the recommendations of Gao et al. to improve the quality, efficiency, and transparency of clustering-based studies (Gao et al., 2023). Unsupervised clustering was performed to identify pneumotypes within ARDS cases. Only DEPs were included for clustering and their log_2_ intensity values were standardized (z-scored) beforehand. To address the curse of dimensionality (Altman and Krzywinski, 2018), we applied principal component analysis (PCA) for dimensionality reduction with the number of principal components (PCs) determined using an ensemble of methods implemented in the R parameters package (version 0.22.0) (Lüdecke et al., 2020). Euclidean distances of PC scores among ARDS cases were then computed and an undirected *k*-nearest neighbors (*k*-NN) graph (with *k* = 12) was constructed. We further built a shared nearest neighbor (SNN) graph (Jarvis and Patrick, 1973) where edge weights were calculated based on Jaccard similarity. Spectral clustering was subsequently performed on this SNN graph using the R igraph package (version 2.1.4) (Csárdi and Nepusz, 2006) based on the random walk normalized Laplacian matrix. The optimal number of clusters *c* was determined by jointly considering the eigengap heuristic and the sample sizes of each cluster. Finally, *k*-means clustering was applied to the selected set of eigenvectors corresponding to the *c* smallest eigenvalues. The derived clusters were visualized by Uniform Manifold Approximation and Projection (UMAP) (McInnes et al., 2020) on the same constructed SNN graph using the R umap package (version 0.2.10.0) (Konopka, 2023).

We assessed the internal coherence of the derived clusters by variance-partitioning analyses of DEP abundances. First, we performed redundancy analysis (RDA) (van den Wollenberg, 1977) to quantify the proportion of variance in the DEPs explained by three sets of variables: cluster labels, baseline variables, and outcome variables. To address the curse of dimensionality (Altman and Krzywinski, 2018), we applied factor analysis of mixed data (FAMD) on the baseline variables using the R FactoMineR package (version 2.12) (Lê et al., 2008) with the number of PCs retained determined by the scree plot. Continuous outcome variables were standardized prior to analysis. RDA was conducted using the R vegan package (version 2.6-10) (Oksanen et al., 2025). In addition, we performed sparse redundancy analysis (sRDA) (Csala et al., 2017) with the R sRDA package (version 1.0.0) (Csala et al., 2017), incorporating variable selection to identify the predictors that best explained the variance in DEPs from a combination of cluster labels, baseline variables, and outcome variables. In this analysis, baseline variables were used directly rather than PCs, and all continuous baseline and outcome variables were standardized. Hyperparameters for the ridge penalty and the number of variables included in the model were tuned using five-fold cross-validation.

### 2.4 Taxonomic biomass proportions and diversity analysis

Microbial proteins were taxonomically annotated using UniProt (release 2025_02) (Consortium, 2024). The complete NCBI taxonomic hierarchy for each identified taxon was retrieved with the R taxize package (version 4.3.0) (Chamberlain and Szöcs, 2013). Taxonomic biomass at the species level and higher taxonomic ranks was calculated by summing the raw intensity values of all detected proteins assigned to each taxon. Relative biomass proportions were subsequently derived based on the contribution of each taxon to the total detected microbial protein intensity, following the approach proposed by Kleiner et al (Kleiner et al., 2017). Median taxonomic biomass proportions were calculated separately for each pneumotype and the control group. Krona visualization and diversity analyses were performed using these median proportions to characterize the distribution of taxon-attributed protein biomass and the diversity of expressed microbial proteins.

Krona plots were generated using Krona Tool (version 2.8.1) (Ondov et al., 2011), in which sector areas correspond to the median biomass proportions across taxonomic hierarchies. For alpha diversity analysis (Whittaker, 1960), indices were selected according to guideline from Cassol et al (Cassol et al., 2025). Species richness was defined as the number of taxa with detected protein biomass signals and represented the richness category. Dominance and evenness were assessed using the Berger-Parker index (Berger and Parker, 1970), calculated as the largest relative biomass proportion among all taxa within a sample. Phylogenetic diversity was quantified using Faith’s phylogenetic diversity (Faith’s PD) (Faith, 1992). Information diversity was evaluated using the Shannon diversity index (Shannon, 1948).

Phylogenetic trees with branch lengths representing estimated divergence times were constructed using TimeTree (version 5) (Kumar et al., 2022). Faith’s PD was calculated using the R picante package (version 1.8.2) (Kembel et al., 2010), whereas Shannon diversity index was computed using R vegan package (version 2.6-10) (Oksanen et al., 2025). Differences in alpha diversity among groups were first assessed using the Kruskal-Wallis *H* test (Kruskal and Wallis, 1952), followed by pairwise comparisons using the Wilcoxon rank-sum test (Wilcoxon, 1945).

For beta diversity analysis (Whittaker, 1960), three complementary dissimilarity metrics were applied. Phylogenetic beta diversity was assessed using the UniFrac (unique fraction metric) distance (Lozupone and Knight, 2005), with both unweighted (Lozupone and Knight, 2005) and weighted (Lozupone et al., 2007) variants calculated using the R package GUniFrac (version 1.9) (Chen et al., 2025a). Aitchison distance (Aitchison, 2018) was used to quantify compositional dissimilarity based on species-level biomass proportions. Principal coordinates analysis (PCoA) (Mead, 1992) was subsequently applied for dimensionality reduction and visualization in two dimensions. Group separation in the reduced ordination space was evaluated using permutational multivariate analysis of variance (PERMANOVA) (Anderson, 2001), implemented in the R vegan package (version 2.6-10) (Oksanen et al., 2025) with 2,000 permutations.

The same Krona visualization and diversity analyses were additionally performed using only species assigned to identified DEPs.

### 2.5 Identification, functional annotation, and clinical correlation analysis of pneumotype-associated DEPs

A univariate logistic regression framework with a one-versus-rest strategy was applied to identify pneumotype-associated DEPs. Each model included a single protein from the identified DEPs as the predictor, without additional covariates. Proteins were considered pneumotype-associated if the association reached statistical significance (*p* < 0.05).

To perform enrichment analysis of pneumotype-associated proteins, protein functional annotation was first conducted using the eggNOG-mapper web server (version 2.1.12) (Cantalapiedra et al., 2021), which assigns annotations based on protein sequences against the eggNOG 5 database (Huerta-Cepas et al., 2019) and maps proteins to corresponding KEGG pathways (Kanehisa et al., 2025) and GO terms (Ashburner et al., 2000; The Gene Ontology Consortium, 2026). Over-representation analysis based on hypergeometric test was then performed separately on KEGG pathways and GO terms using the R clusterProfiler package (version 4.10.1) (Wu et al., 2021) with the background defined as proteins having at least one mapped annotations. Pathways with *p* < 0.05 were considered significantly enriched. Redundancy among enriched GO terms was reduced using the simplify function in the R clusterProfiler package (version 4.10.1) (Wu et al., 2021) with default parameters. Specifically, GO terms with a semantic similarity score based on Wang’s method (Wang et al., 2007) greater than 0.7 were removed, retaining the term with the smallest *p*-value within each similarity group. Subsequently, enriched KEGG pathways and GO terms were further clustered using the R pathfindR package (version 2.4.1) (Ulgen et al., 2019) with default settings. Pairwise similarity between annotation terms was calculated using kappa statistics (McHugh, 2012) based on overlapping hit proteins assigned to each term. Hierarchical clustering was then applied to group annotation terms, with the optimal number of groups determined using the silhouette index (Rousseeuw, 1987). Within each group, the term with the smallest *p*-value was selected as the representative annotation term.

Clinical correlation analysis was performed to evaluate correlations between the abundance of pneumotype-associated DEPs and clinical variables. Since protein abundance represents a continuous variable, correlation methods were selected according to the data type of each clinical variable. Pearson correlation coefficients (Pearson and Galton, 1895) were calculated for continuous clinical variables. For binary variables, point-biserial correlation (Pearson, 1910) was applied. Correlations with ordinal clinical variables were assessed using polyserial correlation (Olsson et al., 1982), implemented using the polyserial function in the R polycor package (version 0.8-2) (Fox, 2025) with maximum likelihood estimation (Cox, 1974). Significant correlations were defined as those with an absolute correlation coefficient greater than 0.35 and a *p*-value < 0.05.

### 2.6 Exploration of clinical characteristics across pneumotypes

We examined differences in clinical baseline and outcome characteristics across pneumotypes identified by unsupervised clustering. For continuous baseline variables and the APACHE III score, distributions were first assessed using the Shapiro-Wilk test (SHAPIRO and WILK, 1965). If all pneumotypes satisfied normality (*p* > 0.05 in each pneumotype), data were compared using Welch’s ANOVA (Welch, 1951). If normality was violated in any pneumotype, data were compared using the Kruskal-Wallis *H* test (Kruskal and Wallis, 1952). Categorical baseline variables, 60-day and 90-day mortalities were compared using Pearson’s χ² test (Pearson, 1900) without Yates’s correction (Yates, 1934), and two-sided Fisher’s exact test (Fisher, 1922) was applied when expected cell counts were < 5. Ordinal variables with more than two levels were compared using Kruskal-Wallis *H* test (Kruskal and Wallis, 1952) to account for their ordinal nature. Survival data were compared with log-rank test (Mantel, 1966). For composite outcomes, ventilator-free days and ICU-free days were compared using the Kruskal-Wallis *H* test (Kruskal and Wallis, 1952) with deaths before day 28 assigned a value of zero. The six organ injury indices were conditional on survival with days counted only while patients were alive. To address death as a competing event, comparisons of these indices were performed among 28-day survivors using the Kruskal-Wallis *H* test (Kruskal and Wallis, 1952). For pairwise comparisons, the Wilcoxon rank-sum test (Wilcoxon, 1945) was applied to variables initially assessed using the Kruskal-Wallis *H* test (Kruskal and Wallis, 1952), while Welch’s *t*-test (WELCH, 1947) was used for variables evaluated with Welch’s ANOVA (Welch, 1951). Significant differences in clinical characteristics were defined as *p* < 0.05.

### 2.7 Identification of pneumotype-associated top-ranked microbial species and pathogens related to 90-day mortality

To examine the association between pneumotype-specific microbial profiles and the primary outcome of 90-day mortality, we focused on the top-ranked microbial species and pathogens within pneumotypes that exhibit 90-day deaths. Species were assigned to a pneumotype-specific rank based on the descending order of the previously calculated median biomass proportions on all detected EV proteins. For each pneumotype, the top 20 ranked species were retained. To focus on species specifically associated with pneumotypes with 90-day deaths, species that appeared among the top 20 in pneumotypes without deaths were excluded from the corresponding top 20 lists of pneumotypes with deaths.

The same ranking strategy was applied to identify top-ranked pathogens, restricting the analysis to species annotated as pathogenic. Pathogen annotations were obtained from three sources: (1) established bacterial pathogens reported by Bartlett et al. (Bartlett et al., 2022); (2) PHI-base version 5.2 (Urban et al., 2025), including only species with *Homo sapiens* as the host; and (3) the Virulence Factor Database (VFDB) (Zhou et al., 2025), updated on November 10, 2025. A species was classified as pathogenic if it was annotated as a pathogen in any one of these sources.

### 2.8 Statistical Analysis

To explore clinical characteristics across pneumotypes, statistical tests were performed in both across all pneumotypes collectively and pairwise comparisons. The choice of statistical test was determined by variable types, including continuous, nominal, and ordinal variables. For continuous variables, the selection of parametric or nonparametric tests was based on an initial assessment of normality. Continuous variables with a normal distribution across all pneumotypes (Shapiro-Wilk test *p* > 0.05 in each pneumotype) are reported as mean ± standard deviation and compared using Welch’s ANOVA. Variables not normally distributed (Shapiro-Wilk test *p* < 0.05 in any pneumotype) are reported as median (25th-75th percentile) and compared using the Kruskal-Wallis *H* test. For nominal categorical variables, statistics are reported as counts (percentage) and compared using Pearson’s χ² test or Fisher’s exact test. Variables involving death as a competing event were analyzed with this consideration incorporated into the statistical framework. *Post hoc* power analysis was conducted based on the observed results, with effect sizes reported for comparisons achieving at least 80% statistical power. Unless otherwise specified, all statistical tests were two-sided, and statistical significance was defined as *p* < 0.05.

## 3. Results

This study aimed to decipher the lung microbial EV protein expression patterns in ARDS patients and to evaluate their predictive roles in the outcomes and severity of the disease (**Figure 1**). Clustering of differentially abundant microbial EV proteins in the BALF samples from ARDS cases and controls was performed to identify potential pneumotypes. To assess the clinical relevance of lung microbial EV protein expression patterns, we explored the association between the identified pneumotypes and the baseline characteristics and outcomes of ARDS.

**Figure 1.**
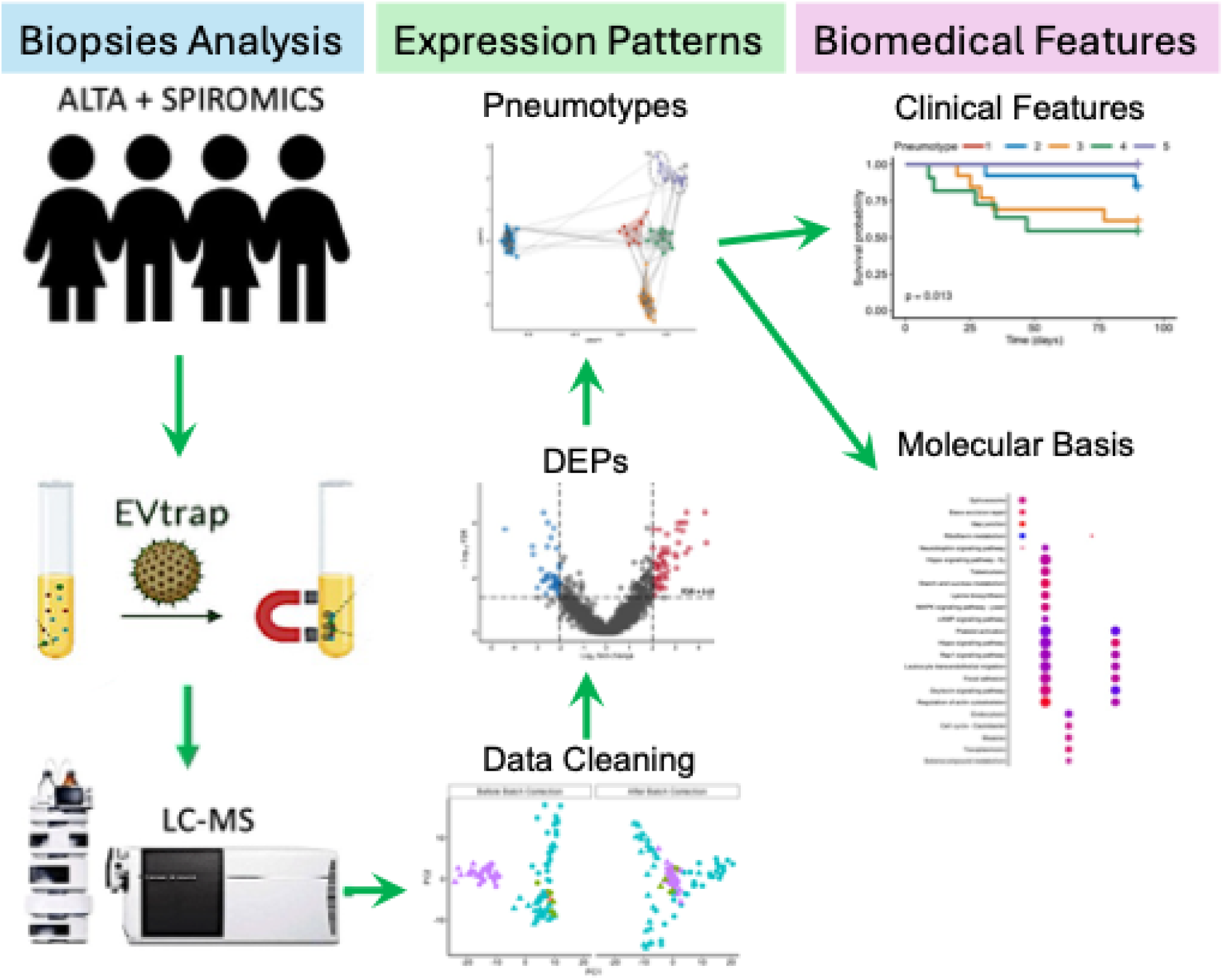
Schematic workflow of the study.

### 3.1 Characteristics of the study cohorts

Demographic and baseline characteristics are summarized in **Table 1** and supplementary Table 1 (**Table S1)**, with no statistically significant differences observed between the ARDS and healthy control groups (all *p* > 0.05). For ARDS cases, a total of 59 baseline variables were collected across nine categories: demographics, ARDS severity, ARDS etiology, comorbidities, medications administered in the last 24 hours, alcohol and smoking behavior, baseline vital signs, baseline laboratory tests, and baseline ventilator parameters. In addition, 12 outcome variables were recorded, including three primary outcomes (overall survival days, 60-day and 90-day death rates) and nine secondary outcomes related to ICU indices and organ failures.

**Table 1.** Demographics and baseline characteristics of ARDS cases and controls. Continuous variables are reported as mean ± standard deviation and compared with the Wilcoxon rank-sum test. Categorical variables are reported as counts (percentage) and compared with the χ² test. Missing counts and percentages are reported only when present within a specific group; otherwise, the cells are left blank. Abbreviation: ARDS, acute respiratory distress syndrome; BMI, body mass index.

| Characteristic | ARDS (n = 57) |  | Control (n = 49) |  | *p*-value |
|---|---|---|---|---|---|
|  | Observed | Missing | Observed | Missing |  |
| Age (years) | 49.40 ± 17.31 |  | 55.41 ± 9.28 |  | 0.060 |
| Female | 24 (42%) |  | 29 (59%) |  | 0.080 |
| Race non-White | 14 (25%) | 1 (2%) | 17 (35%) | 1 (2%) | 0.247 |
| Ethnic Hispanic or Latino | 5 (9%) |  | 6 (12%) |  | 0.559 |
| BMI (kg/m²) | 27.27 ± 6.56 |  | 28.39 ± 4.87 |  | 0.219 |
| Current smoker | 20 (40%) | 7 (12%) | 18 (37%) |  | 0.738 |
| Smoking, pack-years | 20.96 ± 25.91 | 9 (16%) | 26.56 ± 23.94 | 1 (2%) | 0.135 |

### 3.2 Metaproteomic data preprocessing and differential analysis

A total of 1,669 microbial proteins were identified with a mean of 1,119 proteins detected per sample. Of these, 373 proteins (22.35%) were excluded due to a missing rate exceeding 50% in either the ARDS or control group (**Fig. S1a**), leaving 1,296 proteins for subsequent analysis. Following variance-stabilizing normalization (**Fig. S1b**), we reassessed the distribution of missing values. As shown in **Fig. S1c**, samples with missing values exhibited lower intensity profiles compared to those without missing values, consistent with a typical pattern in proteomics data where missingness arises from proteins falling below or near the detection limit. This observation supports the use of a left-censored imputation strategy in our study. After imputation, an increase in the number of detected low-abundance proteins was observed (**Fig. S1d**), confirming the effectiveness of the imputation approach. The primary potential batch factor was the sequencing date of BALF samples. Boxplots of the median protein intensities across sequencing batches did not reveal systematic signal drift (**Fig. S2a**). However, the PCA plot of samples colored by sequencing date indicated a noticeable batch effect in the overall protein abundance profiles (**Fig. S2b**, left). After batch effect adjustment using the ComBat algorithm, samples from different batches showed improved mixing in PCA space (**Fig. S2b**, right), and the difference between within-batch and between-batch correlation distributions was reduced (**Fig. S2c**), indicating effective batch adjustment. After metaproteomic data preprocessing, 1,296 microbial proteins were retained with a median pooled standard deviation of 3.24 across log₂-transformed protein intensities.

Based on a fold change threshold of 4 (corresponding to Cohen’s *d* effect size of 0.62, and *post hoc* statistical power of 82.99% using Wilcoxon rank-sum test), we identified 90 significant DEPs (FDR < 0.05) in lung microbial EVs, including 53 proteins (58.89%) with higher abundance in ARDS cases and 37 proteins (41.11%) with lower abundance compared with healthy controls (**Fig. 2a**, **Table S2**). The abundance of both upregulated and downregulated proteins was clustered (**Fig. 2b**). A PCA plot of the first two components based on these DEPs showed a moderate separation between ARDS and control groups (**Fig. 2c**).

**Figure 2.**
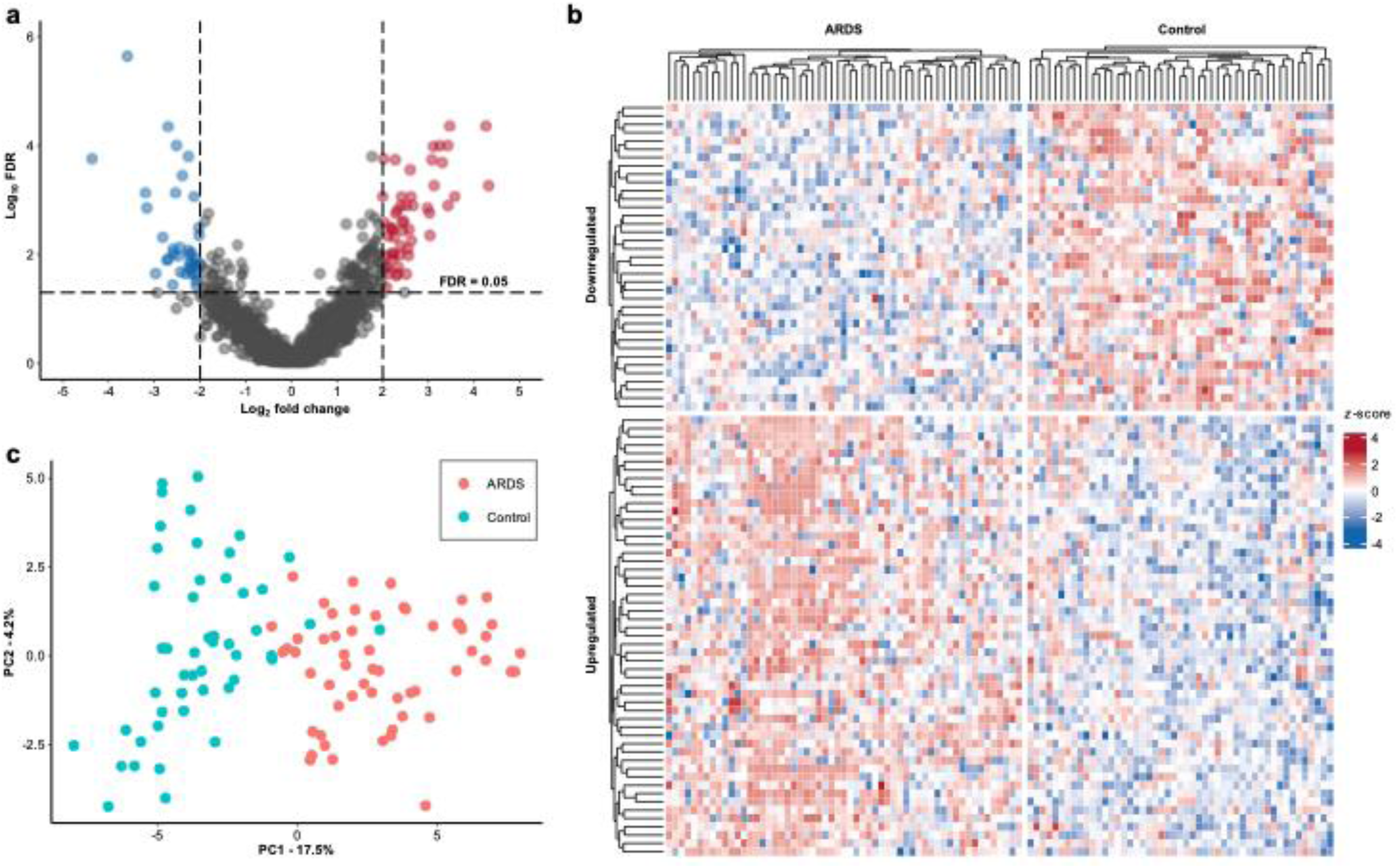
Identification of differentially expressed EV proteins. Differential analysis of microbial EV proteins between 57 ARDS cases and 49 controls using Wilcoxon rank-sum test. (a) Volcano plot. The significance thresholds used to define differentially expressed proteins (DEPs): fold change > 4 and false discovery rate (FDR) < 0.05. (b) Heatmap. (c) Principal component analysis (PCA).

### 3.3 Spectral clustering identified five pneumotypes

A shared nearest neighbor (SNN) graph was constructed using the first three principal components (PCs) of the 90 DEP log_2_ intensities, with the number of PCs suggested by the R package parameters. The eigengap heuristic indicated six clusters from spectral clustering (as shown by the dashed horizontal red line; **Fig. 3a**). However, two closely related clusters contained only a few patients (n = 4 and n = 6, respectively; **Fig. 3b**), and these would merge into a single cluster if the number of clusters was set to five. Therefore, these two small clusters (termed 5A and 5B) were combined to improve the power of sample size. The resulted five clusters were well separated in the UMAP embedding (**Fig. 3b**). To validate cluster coherence, redundancy analysis (RDA) and sparse RDA (sRDA) were performed. RDA indicated that cluster membership explained a substantial portion of variation in DEP abundances (**Fig. 3c**). The total adjusted R² explained by RDA was 10.8%, with residual (unexplained) variance accounting for 89.2%. Cluster labels alone explained 11.5% of the variance, whereas baseline variables contributed minimally (1.2%), and outcome variables explained none (−0.8%). After controlling for baseline and outcome variables, the unique contribution of cluster labels remained significant (adjusted R² = 11.9%; F = 2.23, *p* = 0.001, based on 1,000 permutations under a reduced model). For sRDA, the optimal hyperparameters were identified as a ridge penalty of 0.6 and seven variables were retained in the model. The sRDA revealed a latent component in the variable set that was strongly correlated with a corresponding latent component in the DEP abundance matrix (*r* = 0.919). The selected seven variables and the top ten proteins with the highest absolute weights are shown in **Fig. 3d**. Among the seven variables, two corresponded to cluster labels with relatively large weights (0.67 and −0.28, respectively; **Fig. 3e**), while the remaining five variables had absolute weights < 0.1. Both RDA and sRDA showed that cluster labels explained a greater proportion of variation in DEP abundances than baseline or outcome variables, consistent with their derivation from the DEP matrix. In summary, these five clusters, denoted as pneumotypes P1 to P5, capture intrinsic microbial proteomic signatures.

**Figure 3.**
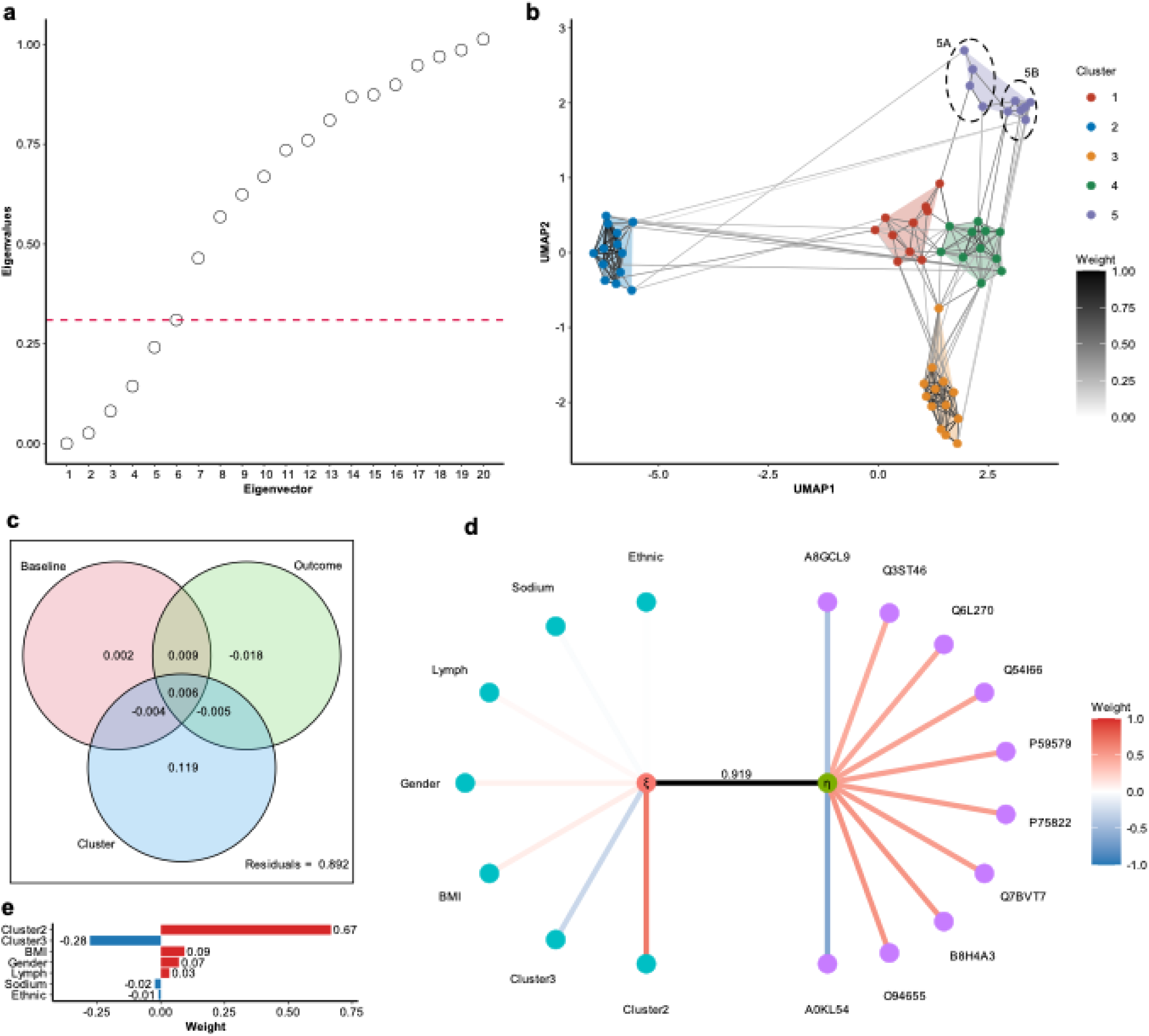
Identification of pneumotypes. (a) Scatter plot of the top 20 eigenvectors with the smallest eigenvalues. The red dashed horizontal line highlights the largest eigengap, corresponding to six clusters. (b) Uniform manifold approximation and projection (UMAP) visualization of the five clusters. Combined new cluster 5 corresponds to original clusters 5 (n = 4) and 6 (n = 6). The UMAP was generated from the same shared nearest neighbor graph used for clustering. (c) Distribution of variance in DEP log_2_ intensities. Values represent adjusted R^2^. RDA, redundancy analysis. (d) Results of sparse RDA showing the seven selected variables and the top ten proteins with the highest absolute weights. Datasets comprising baseline, outcome variables, cluster labels, and the DEP log_2_ intensity matrix are jointly modeled through latent components, revealing strong correlations between the two datasets. (e) Bar plot showing the weights of the seven variables selected in sparse RDA.

### 3.4 Functional enrichment pneumotypes-associated DEPs and their associations with clinical measures

We identified proteins within DEPs that were significantly upregulated or downregulated in one pneumotype compared with the others and defined these as pneumotype-associated DEPs, representing the characteristic protein expression profiles of each pneumotype. Out of the 90 DEPs, univariate logistic regression identified 14, 34, 22, 7, and 17 pneumotype-associated microbial proteins for pneumotypes P1 to P5, respectively (**Table S3**), with minimal overlap among pneumotypes (**Fig. S3**). Functional enrichment analysis of these pneumotype-associated microbial proteins revealed 0, 0, 9, 3, and 1 significantly enriched KEGG pathways, as well as 6, 1, 152, 46, and 30 GO terms. Following redundancy reduction using semantic similarity filtering, 3, 1, 55, 8, and 16 nonredundant GO terms remained for downstream analysis. Subsequent hierarchical clustering of enriched KEGG pathways and GO terms identified 2, 1, 8, 4, and 7 functional annotation groups for P1 to P5, respectively (**Fig. 4a**, **Table S4**). P1’s primary enrichment was “responses to metal ions” and “sulfur compound metabolic process”, indicating a more focused stress-response profile. P2 was enriched only for heterocycle biosynthetic process. P3 was primarily enriched for vesicle-mediated transport, proteolysis, and cellular component disassembly, indicating active protein trafficking and stress adaptation. P4 was dominated by metabolic and biosynthetic processes and Hippo signaling pathway, reflecting enhanced microbial biosynthetic activity. P5 was mainly enriched for organic acid biosynthetic processes, accompanied by proteolysis, suggesting metabolically active microbial functions.

**Figure 4.**
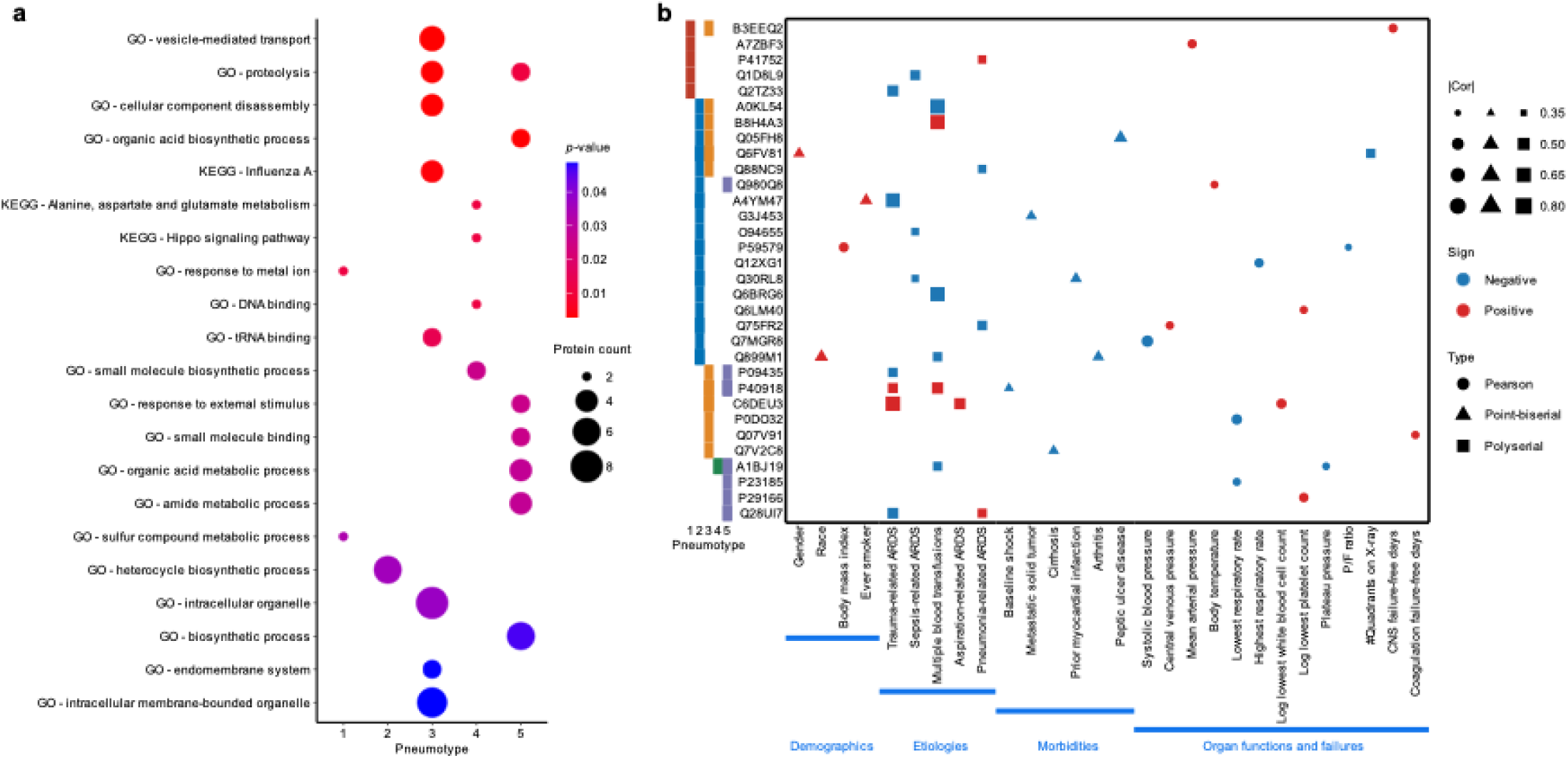
Functional enrichment and clinical correlation analysis of pneumotype-associated differentially expressed proteins (DEPs). (a) Functional enrichment annotation groups obtained by hierarchical clustering of enriched GO and KEGG terms derived from pneumotype-associated DEPs. (b) Significant correlations between pneumotype-associated DEPs and clinical variables, defined by an absolute correlation coefficient greater than 0.35 and a *p*-value < 0.05.

Clinical correlation analysis identified significant associations between 32 pneumotype-associated microbial proteins and 28 clinical variables by an absolute correlation coefficient greater than 0.35 and a *p*-value < 0.05 (**Fig. 4b**). Correlation profiling revealed pneumotype-specific correlation patterns between microbial proteins and host clinical characteristics. Pneumotype P1 showed predominant associations with ARDS etiological factors, along with a positive correlation with central nervous system (CNS) failure-free days. P2 exhibited a broader spectrum of correlations spanning ARDS etiology, comorbidities, demographic characteristics, baseline vital signs, and laboratory indices, characterized by a mixture of positive and negative correlations across clinical variable categories. P3 displayed several associations with ARDS etiological factors, and negative correlation with cirrhosis. In contrast, P4 demonstrated a single correlation pattern, characterized by negative associations involving transfusion and the ventilation-related variable plateau pressure. P5 showed associations primarily related to ARDS etiological factors, as well as respiratory rate and platelet count. Overall, both the correlation signs and clinical variable categories of significant associations varied across pneumotypes, indicating heterogeneous host-microbial interaction patterns.

### 3.5 Divergent clinical characteristics of pneumotype-associated patients

For analyses involving five pneumotypes, with α = 0.05 and 80% statistical power, the estimated effect sizes were 0.48 for the one-way ANOVA and 0.57 for the χ² test based on a 5×2 contingency table. The most pronounced difference in clinical measures among patients of the five pneumotypes was overall mortality rates in 60- and 90-days post admission. In 60 days, P4 patients showed the highest mortality rate (45.45%). P3 and P2 patients had moderate and mildly elevated mortality rates (30.77% and 7.69% for P3 and P2, respectively; *p* = 0.011; **Fig. 5a**, **Fig. 5b**, **Table 2**, **Table S5**). In contrast, all P1 and P5 patients survived. Similarly, significant differences in overall mortality rates were observed in 90 days between pneumotypes, leading by P4 patients (45.45%) followed by P3 (38.46%) and P2 subgroups (15.38%) (*p* = 0.013; **Fig. 5c**, **Table 2**, **Table S5**). These results suggest that most of deaths took place within 60 days with a slight increase in the following 30 days. In addition, increased APACHE score was observed in P3 patients (*p* = 0.016 vs P1 group; **Fig. 5d**, **Table 2**, **Table S5**), while the P/F ratio did not differ among groups with deceased patients (**Fig. 5e**). Regarding secondary outcomes, after excluding the five subjects who died within 28 days to account for death as a competing event, no statistical significance in organ failures was found among P1 and P5 survivors (all *p* > 0.05, **Table S6**). Notably, CNS failure-free days in P2 and P3 patients were reduced to 41% and 27%, respectively, compared with P1 survivors (**Table S6**). Correspondingly, significant reductions in organ failure-free days were observed in these patients (63% and 46%, respectively, for P2 and P3, **Table S6**). These results suggest that the causes of death in P2 and P3 patients could be different from those in P4 patients. Further, to address this possibility, we compared potential differences in etiology and medications. P2 patients had significantly less sepsis (*p* < 0.05 vs P1, **Fig. 5f**, **Table 2**, **Table S7**), while P3 and P4 patients tended to have shock and hypertension (*p* < 0.05, **Fig. 5g-5h**). Differences in interventions and key physiological measures were also compared between groups. P3 and P4 patients who were treated with vasopressors and inotropes within the last 24 hours were more numerous than in other groups (*p* < 0.05, **Fig. 5i**); P3 patients had higher positive end-expiratory pressure (PEEP) than other groups (*p* < 0.05, **Fig. 5j**). Additionally, reductions in hematocrit level (**Fig. 5k**) and body temperature (**Fig. 5l**) were measured (*p* < 0.05 vs P1). Taken together with the correlation between pneumotype-associated DEPS and clinical variables (**Fig 4b**), lung microbial expression patterns (P1-P5) link to patients’ clinical characteristics.

**Figure 5.**
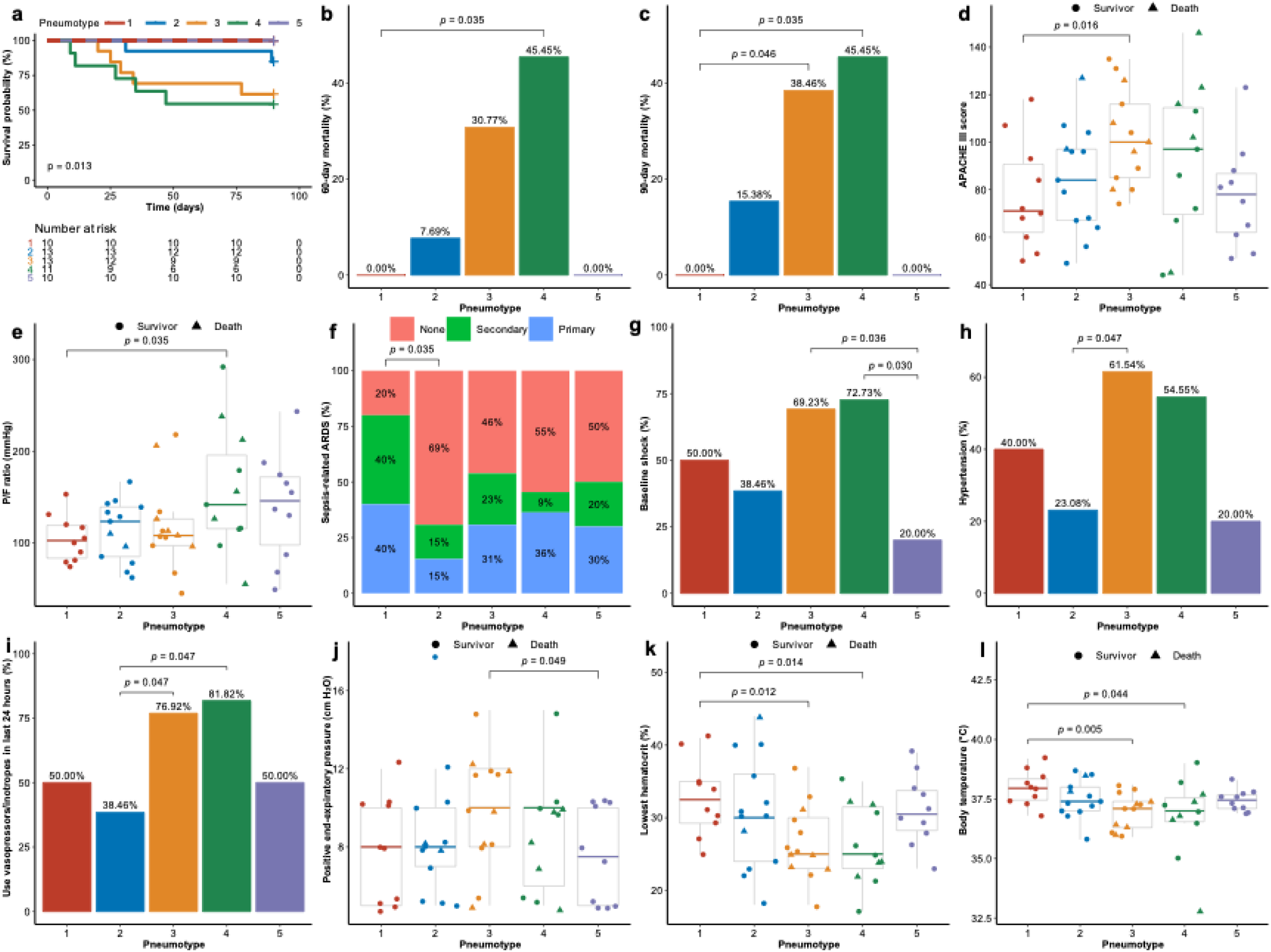
Clinical characteristics of five pneumotypes. (a) Kaplan-Meier survival curves during a 90-day follow-up. Log-rank test. (b-l) Clinical measures differentiating pneumotypes. PaO₂/FiO₂ (P/F) ratio, positive end-expiratory pressure (PEEP). Fisher’s exact test for b, c, g, and i. Welch’s *t*-test for d, e, k, and l. Wilcoxon rank-sum test for f and j. χ² test for h and i.

**Table 2.** Comparison of clinical characteristics between pneumotypes. * Ordinal variables were analyzed using the Kruskal-Wallis *H* test. # *p* < 0.05 for the comparison vs P1 or P5. & p < 0.05 vs P2.

| Characteristic | P1<br>(n = 10) | P2<br>(n = 13) | P3<br>(n = 13) | P4<br>(n = 11) | P5<br>(n = 10) | *p*-value |
|---|---|---|---|---|---|---|
| 60-day mortality | 0 (0.00%) | 1 (7.69%) | 4 (30.77%)<sup>#</sup> | 5 (45.45%)<sup>#</sup> | 0 (0.00%) | 0.011 |
| 90-day mortality | 0 (0.00%) | 2 (15.38%) | 5 (38.46%) | 5 (45.45%)<sup>#</sup> | 0 (0.00%) | 0.013 |
| APACHE III | 77.49 ± 22.69 | 84.15 ± 22.76 | 101.85 ± 20.40<sup>#</sup> | 91.91 ± 32.55 | 77.50 ± 21.77 | 0.065 |
| P/F ratio | 104.97 ± 25.39 | 113.75 ± 33.20 | 118.14 ± 47.73 | 157.13 ± 68.54<sup>#</sup> | 139.58 ± 59.01 | 0.173 |
| Sepsis-related ARDS |  |  |  |  |  | 0.354<sup>*</sup> |
| None | 2 (20%) | 9 (69%)<sup>#</sup> | 6 (46%) | 6 (55%) | 5 (50%) |  |
| Primary | 4 (40%) | 2 (15%) | 3 (23%) | 1 (9%) | 2 (20%) |  |
| Secondary | 4 (40%) | 2 (15%) | 4 (31%) | 4 (36%) | 3 (30%) |  |
| Baseline shock | 5 (50%) | 5 (38%) | 9 (69%)<sup>#</sup> | 8 (73%)<sup>#</sup> | 2 (20%) | 0.083 |
| Hypertension | 4 (40%) | 3 (23%) | 8 (62%)<sup>#</sup> | 6 (55%) | 2 (20%) | 0.171 |
| Vasopressors or inotropes | 5 (50%) | 5 (38%) | 10 (77%)<sup>&</sup> | 9 (82%)<sup>&</sup> | 5 (50%) | 0.137 |
| PEEP (cm H<sub>2</sub>O) | 8.00 (5.00 - 10.00) | 8.00 (7.00 - 10.00) | 10.00 (8.00 - 12.00)<sup>#</sup> | 10.00 (5.00 - 10.00) | 7.50 (5.00 - 10.00) | 0.282 |
| Lowest hematocrit (%) | 32.70 ± 5.27 | 30.62 ± 7.79 | 26.62 ± 5.09<sup>#</sup> | 26.28 ± 5.53<sup>#</sup> | 31.00 ± 4.90 | 0.041 |
| Body temperature (°C) | 37.96 ± 0.72 | 37.51 ± 0.81 | 36.98 ± 0.76<sup>#</sup> | 36.75 ± 1.66<sup>#</sup> | 37.43 ± 0.45 | 0.049 |

### 3.6 Pneumotype P2 exhibits distinct microbial biomass diversity and compositional structure

The total of 1,669 detected microbial proteins corresponded to 626 distinct species following taxonomic annotation. Krona visualization of pneumotypes P1 to P5 and the control group demonstrated broadly similar taxonomic compositions at the domain level, with Bacteria representing the predominant component across all groups (**Fig. 6a-6f**). A phylogenetic tree constructed from 1,501 mapped taxa, with branch lengths scaled according to estimated divergence times, was used for downstream phylogenetic diversity analyses (**Fig. S4**).

**Figure 6.**
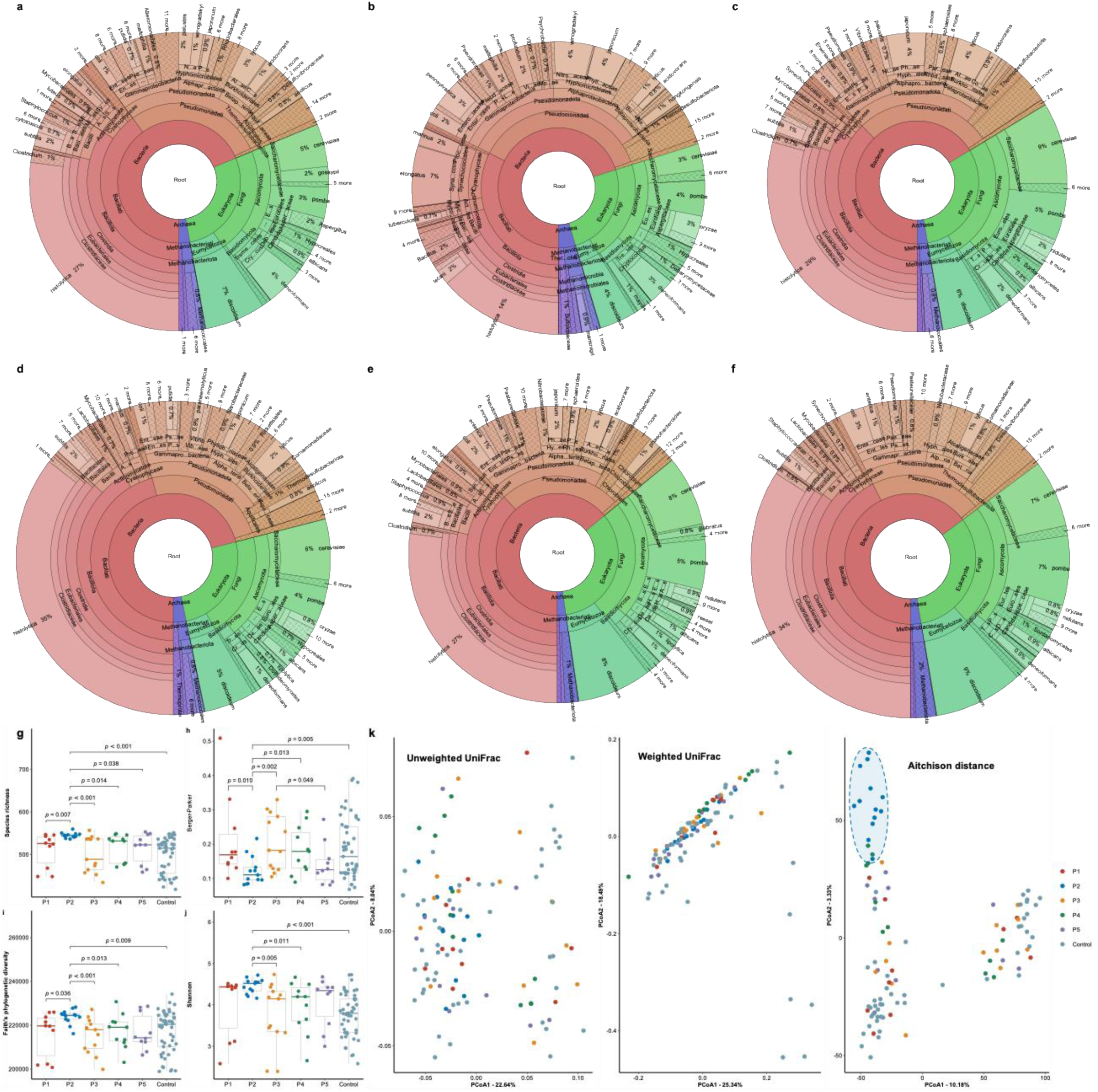
Characteristics of species assigned to all detected microbial proteins across pneumotypes and the control group. (a-f) Krona plots illustrating the taxonomic composition of species corresponding to all detected microbial proteins across pneumotypes P1-P5 and controls. Proportions represent the relative taxon-attributed protein biomass contributions, calculated from summed protein intensities. Alpha diversity indices derived from the same species-level biomass proportions include (g) species richness, (h) Berger-Parker index, (i) Faith’s phylogenetic diversity, and (j) Shannon diversity index. *P*-values were obtained using the Wilcoxon rank-sum test. (k) Beta diversity analysis, including two-dimensional principal coordinates analysis (PCoA) ordination based on unweighted (left) and weighted (middle) UniFrac (unique fraction metric) distances and Aitchison distance (right). The clustered P2 samples in the Aitchison distance-based two-dimensional PCoA visualization are highlighted by a dashed ellipse.

Alpha diversity analysis revealed that P2 exhibited distinct diversity characteristics compared with the other four pneumotypes and the control group. Specifically, P2 showed higher species richness, Faith’s PD, and Shannon diversity index values, together with a lower Berger-Parker dominance index (**Fig. 6g-6j**, **Table S8**), indicating a more heterogeneous and less dominance-driven metaproteomic biomass composition relative to other groups.

Consistent with these findings, beta diversity analysis further demonstrated compositional differences among groups. Two-dimensional ordination based on Aitchison distance showed clear clustering of P2 samples, distinctly separating them from other pneumotypes and controls (**Fig. 6k**). Group-level differences in metaproteomic composition were statistically supported by PERMANOVA analysis (*p* = 0.016).

The same Krona visualization and diversity analyses were additionally performed using only species assigned to identified DEPs. The total of 90 DEPs corresponded to 77 distinct species, and a phylogenetic tree constructed from 65 mapped taxa was used for phylogenetic diversity analysis (**Fig. S5**). Krona visualization across pneumotypes P1-P4 demonstrated that Bacteria remained the predominant domain; however, in P5, Eukaryota exceeded Bacteria as the dominant component (**Fig. S6a-6f**). Alpha diversity analysis based on DEPs showed patterns consistent with those observed for all detected microbial proteins. P2 continued to exhibit higher species richness, Faith’s PD, and Shannon diversity index values, together with a lower Berger-Parker index (**Fig. S6f-6i**; **Table S9**), indicating a more heterogeneous protein biomass composition relative to other pneumotypes. In contrast, beta diversity analysis did not demonstrate significant compositional differences among pneumotypes (**Fig. S6j**).

### 3.7 Divergent microbial features of pneumotypes

To compare the differences in lung microbiota among the top 20 most abundant taxa between survivors and deceased patients, we excluded the annotated microbial species which shared with P1 and P5 subtypes for survivors from P2, P3, and P4 subgroups. Eight, three, and five top-ranked species were retained in P2, P3, and P4, respectively, showing only limited overlap among pneumotypes (**Fig. 7a**). *Aspergillus oryzae* was shared across all three deceased pneumotypes, whereas *Prochlorococcus marinus* was shared between P2 and P4, and *Photobacterium profundum* was shared by P2 and P3. On the other hand, five, one, and three species were specific for P2, P3, and P4, respectively. Among them, P2 and P4 had two different annotated pathogens, *Clostridium tetani* and *Laribacter hongkongensis* for P2, and *Vibrio parahaemolyticus* and *Vibrio cholerae* for P4. In contrast, *Pyricularia oryzae* was the only specific microbe for P3.

**Figure 7.**
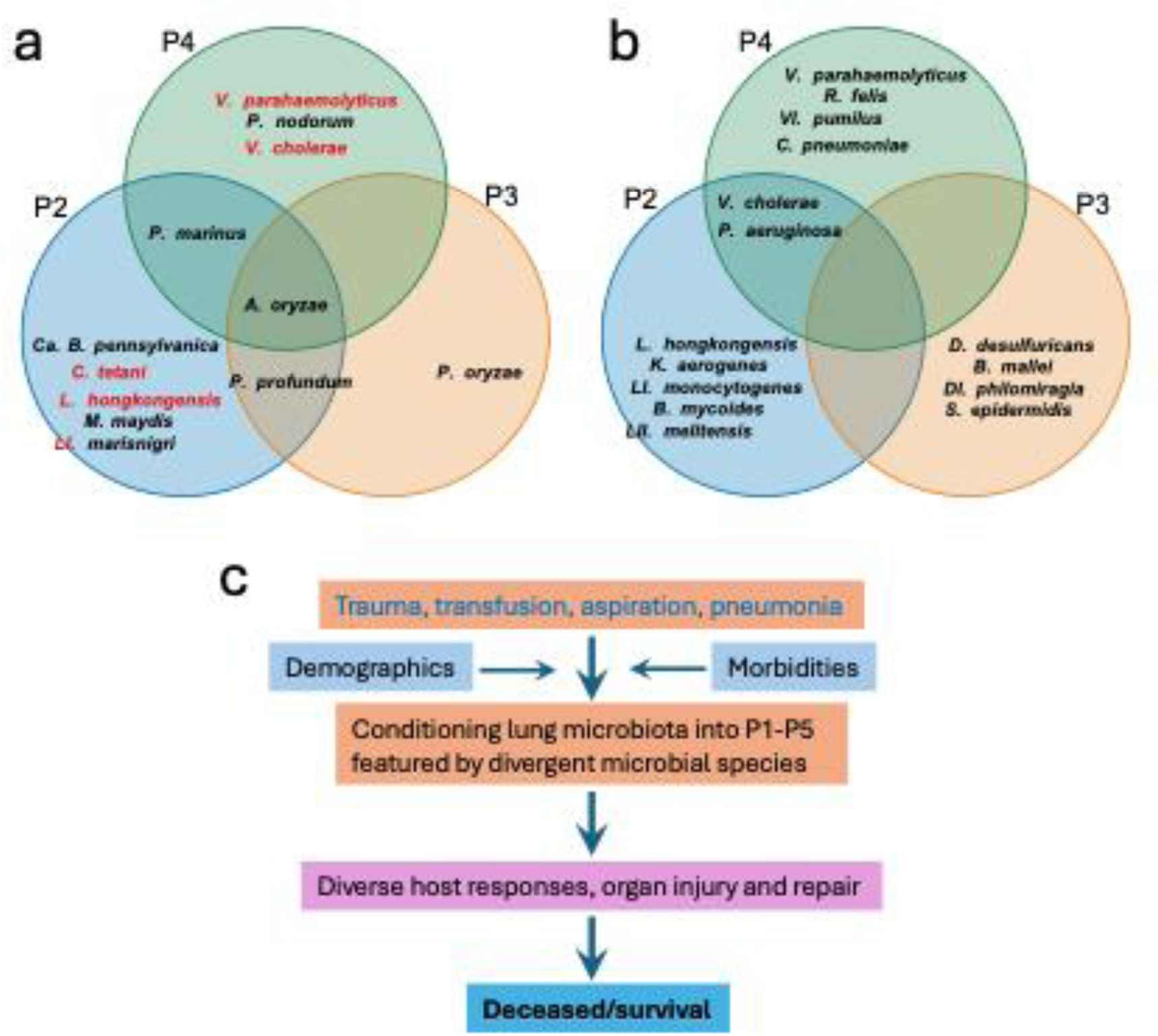
Venn diagrams of top ranked species and pathogens in pneumotypes 2, 3, and 4. Species were ranked by median species-level biomass contribution proportions across samples within individual pneumotype. Species shared with those of pneumotypes 1 or 5 were excluded. (a) Distribution of the top 20 microbial species across the three pneumotypes with deaths, as visualized by a Venn diagram. Species in red font were annotated as pathogens. (b) Venn diagram of the top 20 pathogens across the three pneumotypes. (c) Schematic working model.

Moreover, we annotated and compared top-ranked pathogens between deceased P2-P4 subgroups (**Fig.7b**). After excluding shared pathogens with survived P1 and P5 patients, seven, four, and six pathogens had remained for P2, P3, and P4, respectively. Only two pathogens were shared by P2 and P4: *Vibrio cholerae* and *Pseudomonas aeruginosa*. P2 was characterized by five pneumotype-specific pathogens, including less invasive *Laribacter hongkongensis, Klebsiella aerogenes, Listeria monocytogenes, Bacillus mycoides, and Brucella melitensis*. P3 proteins enriched four pathogens: *Desulfovibrio desulfuricans, Burkholderia mallei, Francisella philomiragia, and Staphylococcus epidermidis.* P4 was specifically featured by four pathogens: invasive *Chlamydia pneumoniae, Vibrio parahaemolyticus, Rickettsia felis, and Bacillus pumilus*.

The schematic working model has been proposed based on this study (**Fig.7c**)

## Discussion

Profiling microbial EV protein expression patterns proteome wide in the lower respiratory tract in ARDS patients, to the best our knowledge, is novel. The bronchoalveolar lavage fluid (BALF) collected using NIH-standard protocol and the well-developed design of ARDSNets and SPIROMIC clinical trials warrant the quality of this study. In parallel analysis of BALF samples collected from both healthy controls and ARDS patients using high-throughput sensitive nano-LC/MS eliminated potential biases. EV proteins demonstrate a long-term stability in contrast to EV-free counterparts, and EV proteomics has advantages over the traditional proteomics of whole plasma proteins. Compared with microbial genomics, EV proteomics is ultimately impossible to be contaminated by environmental microbes. We identified five microbial EV expression patterns in the lower respiratory tract in ARDS patients. These EV pneumotypes are featured by diverse biological processes/functions, associations with different clinical measures, and top-ranked pathogens. For the first time, we have demonstrated that microbial EV proteins can be applied to characterize the changes of lung microbiota in ARDS patients.

Our study identified reasonable microbial EV proteins in BALF samples of ARDS patients. The finding of these upregulated and downregulated DEPs supports previous studies in lung microbiota that pulmonary diseases affecting microbial species. The clustering analysis of DEPs (heatmap) confirms a heterogeneous distribution of these proteins, which is further substantiated by the diffuse locations on the PCA plot. Intriguingly, these DEP proteins significantly separate ARDS from healthy controls, indicating a potential of most of the DEPs may serve as diagnosis biomarkers for the disease. The differences detected in the EV protein expression between controls and ARDS are most unlikely induced by insignificant differences in demographics and baseline feature.

Spectral clustering, a graph-based clustering approach, has demonstrated robust performance in comprehensive benchmarking studies (von Luxburg, 2007; Rodriguez et al., 2019), and has been successfully applied to identify biologically relevant endotypes, including metabo-endotypes in asthma (Kelly et al., 2022). To preserve clustering validity, we avoided any information leakage from clinical variables into the clustering process, as emphasized by Altman et al (Altman and Krzywinski, 2017) and Kelly et al (Kelly et al., 2022). Appropriate dimensionality reduction was applied to mitigate the curse of dimensionality, and a shared nearest-neighbor graph was constructed to enhance clustering stability and robustness. Variation-partitioning analysis further confirmed that clinical variables were not incorporated into cluster construction. Therefore, the patterns observed between clinical characteristics and the resulted unsupervised clusters support the biological and clinical interpretability of the identified pneumotypes (Kelly et al., 2022).

To enrich the EV proteins with GO and KEGG databases, inherent biological processes are characterized for all five identified pneumotypes. Comprehensive host functions have been initiated by these EV proteins. Some of these microbial EV proteins of P3 triggered the host immune responses to infections and subsequent alterations in lung microbiota, including biofilm matrix, response to antibiotics, defense to other organisms, protein targeting ER and mitochondria, and estrogen and MAPK signaling pathways. By comparison, it showed fewer biological functions are targeted by P2 and P3 proteins. In survival patients, the targeted host functions are predominately limited to metabolic processes. The functional differences targeted by pneumotype-associated DEPs may partially explain divergent mortality and other associated clinical variables.

Individual microbial EV proteins were associated with etiologies and organ functions. These results suggest that the lower airway microbiota may be altered by different causes and invaded pathogens. Subsequently, organ dysfunction and systemic abnormality may occur following altered lung microbial community. Only one microbial protein, in sharp contrast, was associated with each measure of demographics and morbidity, indicating that demographics and pre-existing conditions may not serve as confounders for the links between these correlated microbial EV proteins and etiology/organ injury. The identified pneumotypes exhibit significant difference in clinical measures, in particular, between the deceased and survived groups. Among them, P3 patients share more clinical dysfunctions with P4 group, including progressive deaths within 90 days, hypertension history, shock and treatments, greater ICU severity, stronger ventilation support, reduced hematocrit level and body temperature. All these clinical disorders are related to impaired cardiovascular system. Survived P1 patients are characterized by the highest rate of sepsis-induced ARDS. In contrast to P3 and P4 patients, less dysfunctional cardiovascular system may benefit the survival in P1 patients. P2 patients have the lowest sepsis-induced ARDS combined with lesser ICU support compared with P3 and P4 groups, which may contribute to the reduced mortality. P5 group has the lowest incidence of shock and pre-existing hypertension. The survival of P5 patients may be attributed to better cardiovascular system. Obviously, these five pneumotypes have distinct clinical features.

Diverse expression patterns may exist at the species level, as shown by alpha and beta diversities. Intriguingly, P2 standouts for both richness and community composition. The pneumotype-specific, top-ranked microbes and pathogens could be explained at least partially by the different etiologies, morbidities, and organ dysfunction (**Fig. 7C**). Our study provides the evidence to support the following model: various etiologies condition microbiota in the lower airway track in ARDS patients, and EV proteins were released into BAL fluid with five expression patterns. These five pneumotypes possess featured microbes and pathogens. Different host responses are triggered by the subgroup-specific microbial EV proteins subsequently, including immune responses and cell/tissue injury and repair. Clinically, clustered groups represent various organ dysfunction and survival rates.

This study has some limitations. ARDSNet clinical trials were designed to test the benefits of therapeutics on ARDS patients. All participants were severe patients randomly enrolled to receive a placebo or a test compound. To have healthy controls, we included BALF samples collected from SPIROMIC trial that was designed for COPD. All healthy controls did not have COPD and ARDS. Both clinical trials were supported by the same institute NHLBI and applied to the same procedure to sample BAL biopsies. Different microbial species release EV unequally in diseased airways. It is possible that EV proteome might not have equal sensitivity to all components of lung microbiota. Causal inference and validation in preclinical models may identify the pneumotype-specific pathogens. The analysis of phenotype-based pneumotypes is limited by the size of cohort.

Our study has identified five new pneumotypes by clustering microbial EV proteins proteome wide in the lower respiratory tract in ARDS patients. These pneumotypes are characterized by divergent microbiota associated with different clinical features, triggered host responses, and potential pathogenicity. The comparison between EV metaproteomics and 16S RNA assay may provide additional evidence for the applicability of microbial EV metaproteome in the study of lung microbiota. These novel observations await further studies to validate the findings, particularly, longitudinal prospective cohort studies.

## Author contributions

HLJ conceived and designed the study. NL, HLJ, and RZZ contributed to the writing of the manuscript. NL, RZZ, AB, and HLJ analyzed the data and performed the statistical analysis. JZ, DS, and EG edited manuscript. All authors reviewed the manuscript and provided critical comments.

## Acknowledgments

This work was supported by NIH NHLBI grant HL87017 (to HLJ) and institutional funds (to HLJ).

## Ethics approval and consent to participate

This study was approved by an exempted Institutional Review Board approval at Loyola University Chicago.

## Disclosure for Biopsies

This study was prepared using the research materials of ARDSNet and SPIROMICS clinical trials obtained from the NHLBI Biologic Specimen and Data Repository Information Coordinating Center (BioLINCC) and does not necessarily reflect the opinions or views of these clinical trial studies or the NHLBI.

