## Supplementary Files for "Deciphering lung microbial EV pneumotypes in acute respiratory distress syndrome proteome wide"

**Funding information**

NIH HL134828, LUC LU103613.

**Supplementary Methods**

### 1. Preprocessing of Clinical Variables from Clinical Trials

**Data extraction.** Clinical variables were extracted from the respective trial documentation. For the SPIROMICS cohort, only baseline demographic variables were extracted, including age, sex, race, ethnicity, body mass index (BMI), and smoking status (current smoker and cumulative smoking exposure measured in pack-years). For the ARDSNet cohort, a broader set of variables was available, including 68 baseline characteristics and 12 follow-up outcomes.

**Removal of variables with high missingness or no variability**. Within the ARDSNet dataset, baseline variables with high missingness (> 20%) were excluded from further analysis. Specifically, the following five variables were removed due to missing data proportions:

- serum total protein (g/dL): 57.89%
- serum albumin (g/dL): 50.88%
- serum phosphorus (mg/dL): 47.37%
- serum magnesium (mEq/L): 45.61%
- highest serum bilirubin (mg/dL): 24.56%

Additionally, four variables exhibiting no variability within the acute respiratory distress syndrome (ARDS) cases (i.e., only one observed level) were removed, including leukemia, hepatic failure with coma or encephalopathy, prior stroke with sequelae, and dementia.
Following this filtering, 59 baseline variables were retained. Outcome variables were unaffected by this step.

**Extreme value clipping and log transformation**. To reduce the influence of extreme outliers, selected variables were winsorized at the 1st and 99th percentiles:

- Clipped at lower bound (1st percentile): serum potassium (mEq/L) and the lowest hematocrit (%)
- Clipped at upper bound (99th percentile): total respiratory rate (breaths/min), total minute ventilation (L/min), hemoglobin (g/dL), serum bicarbonate (mEq/L), highest serum creatinine (mg/dL), heart rate (beats/min), and central venous pressure (mmHg)

To address skewed distributions, serum glucose (mg/dL), the lowest white blood cell count ($mm^3$), the lowest platelet count ($\times 1{,}000/mm^3$), and the highest blood urea nitrogen (mg/dL) were log-transformed.
Outcome variables were not altered during this clipping and transformation process. Clinical baseline variables were then grouped into nine categories, including:

1) Demographics (5 variables)
2) ARDS severity (2 variables)
3) ARDS etiology (6 variables)
4) Comorbidities (14 variables)
5) Medicines in last 24 hours (2 variables)
6) Alcohol & Smoking (4 variables)
7) Baseline vital signs (8 variables)
8) Baseline lab tests (10 variables)
9) Baseline ventilator (8 variables)

And the outcome variables were grouped into two categories: primary and secondary outcomes.

1) Primary outcome - mortality: Three variables were defined, including the number of days to death during follow-up (up to 90 days), and two additional indicator variables representing 60-day and 90-day mortality.

2) Secondary outcomes - intensive care unit (ICU) index and organ failures: ICU index includes 3 variables: ventilator-free days, ICU-free days and APACHE III score, and organ failures up to 28 days includes 6 variables: cardiovascular failure-free days, central nervous system (CNS) failure-free days, coagulation failure-free days, renal failure-free days, hepatic failure-free days, and total days without failure of any of the aforementioned organs.

Ventilator-free days and ICU-free days were recorded with deaths within 28 days assigned a value of zero, thereby accounting for the competing risk of mortality[1]. For the six organ failure indices, the counting of failure-free days was terminated at the time of death. Detailed descriptions of the 59 baseline variables and 12 outcome variables in the ARDS cohort are provided in Supplementary file for Table S1.

**Missing data imputation.** To prepare the data for downstream statistical testing and regression-based modeling, missing values in the 59 baseline variables were imputed using the missForest package in R (version 1.5)[2], which leverages observed values across mixed-type features to iteratively predict missing entries using random forest models.

When imputing the baseline variables, only baseline variables were used for the imputation. Outcome variables were excluded from the imputation process to prevent potential information leakage. Notably, outcome variables were largely completed; only one outcome variable (APACHE III score) had a single missing value, which was imputed using the available baseline variables.

**2. Extracellular Vesicle Protein Extraction from BALF and LC-MS/MS Data Processing**

Extracellular vesicles (EVs) were isolated from bronchoalveolar lavage fluid (BALF) as previously described[3, 4]. Briefly, exosomes contained within EVs were purified, and their protein contents were extracted for liquid chromatography-tandem mass spectrometry (LC-MS/MS) analysis (Thermo Fisher Scientific). Peptide intensities were obtained, and raw mass spectral data were converted into LC-MS and LC-MS/MS spectra using MSConvert in the ProteoWizard suite[3, 4].

**Microbial protein database construction**. The reference database comprised protein sequences from all bacterial, archaeal, fungal, viral, and slime mold (*Dictyostelium discoideum*, *Physarum polycephalum*) species, as well as *Homo sapiens*, downloaded from the Swiss-Prot section of UniProtKB (release 2021_04)[5].

**Peptide identification**. Database searching was performed using Comet (version 2018.01 rev.2)[6]. Default parameters were applied except for the following settings: precursor mass tolerance of 0.1 Da; peptide mass range from 400 to 11,000 Da; enzyme specificity set to Trypsin/P; up to three missed cleavages; a maximum of three variable modifications per peptide; variable modifications of methionine oxidation (+15.9949 Da) and N-terminal acetylation (+42.010565 Da); and a static cysteine carbamidomethylation (+57.0214 Da).

**PSM filtering, protein grouping and quantification**. Peptide–spectrum match (PSM) filtering and label-free quantification were conducted using the Sipros/ProRata workflow[7]. PSMs were filtered at a false discovery rate (FDR) of 0.2% at the PSM level using a target-decoy approach[8]. Protein-level FDR varied across samples, with the highest observed FDR being 1.64%. Prior to quantification, MS1 retention times were aligned using the retention time alignment algorithm described by Katharina et al[9]. Protein abundances were quantified based on the total chromatographic peak area computed by ProRata under default parameters. Within the Sipros/ProRata framework, identified proteins were organized into protein groups, defined as sets of proteins sharing the same identified peptide sequences and therefore indistinguishable at the peptide level. Only protein groups containing at least one unique peptide were retained for downstream analysis. For groups comprising multiple proteins, total abundance was evenly distributed among constituent members to obtain protein-level quantitative estimates. Finally, only proteins mapped to the microbiome were retained for subsequent analyses.

**Taxonomic annotation**. Microbial proteins were taxonomically annotated based on their recognized UniProt accession identifiers using UniProt (release 2025_02)[5]. The complete NCBI taxonomic hierarchy for each taxon was retrieved with the taxize package (version 4.3.0)[10].

#### 3. Metaproteomic Data Preprocessing

The preprocessing of metaproteomic data involved sequential steps of filtering, normalization, imputation, and batch effect correction. First, proteins with excessive missingness were excluded; specifically, any protein with more than 50% missing values in either the ARDS or control group was discarded. Next, variance-stabilizing normalization[11] was applied using the R package vsn (version 3.70.0)[11], following the benchmark established by Välikangas et al[12]. To address missing data, we performed left-censored missing data imputation using the MinProb method from the imputeLCMD package (version 2.1)[13], as recommended by Lin et al[14]. This approach imputes missing values by drawing randomly from a Gaussian distribution where the mean is set to the 1st percentile of the observed protein intensities within each sample. Finally, we assessed potential batch effects using principal component analysis (PCA) and performed intra-batch signal drift inspection. When batch effects were detected, they were adjusted using the empirical Bayes framework implemented in the ComBat algorithm[15] from the sva package (version 3.50.0)[16].

**Table S1**. Clinical characteristics of the ARDS cohort.

| Characteristic | | Missing | ARDS (n = 57) |
|---|---|---|---|
| ***Demographics* (5 variables)** | | | |
| Age (years) | | | 49.40 ± 17.31 |
| Female | | | 24 (42%) |
| Race non-White | | 1 (2%) | 14 (25%) |
| Ethnic Hispanic or Latino | | | 5 (9%) |
| Body mass index (kg/m²) | | | 27.27 ± 6.56 |
| ***ARDS severity* (2 variables)** | | | |
| P/F ratio (mmHg) | | | 126.11 ± 50.77 |
| Number of quadrants with opacities on chest X-ray | | | |
| | 2 | | 8 (14%) |
| | 3 | | 13 (23%) |
| | 4 | | 36 (63%) |
| ***ARDS etiology* (6 variables)** | | | |
| Trauma-related ARDS | | | |
| | None | | 52 (91%) |
| | Primary | | 4 (7%) |
| | Secondary | | 1 (2%) |
| Sepsis-related ARDS | | | |
| | None | | 28 (49%) |
| | Primary | | 17 (30%) |
| | Secondary | | 12 (21%) |
| Multiple blood transfusions | | | |
| | None | | 53 (93%) |
| | Primary | | 2 (4%) |
| | Secondary | | 2 (4%) |
| Aspiration-related ARDS | | | |
| | None | | 38 (67%) |
| | Primary | | 13 (23%) |
| | Secondary | | 6 (10%) |
| Pneumonia-related ARDS | | | |
| | None | | 21 (37%) |
| | Primary | | 21 (37%) |
| | Secondary | | 15 (26%) |
| Other lung injury | | | |

| | | |
|---|---|---|
| None | | 55 (96%) |
| Primary | | 1 (2%) |
| Secondary | | 1 (2%) |
| ***Comorbidities* (14 variables)** | | |
| Baseline shock | | 29 (51%) |
| Chronic dialysis | | 4 (7%) |
| Non-Hodgkin's lymphoma | | 1 (2%) |
| Metastatic solid tumor | | 1 (2%) |
| Immunosuppression | | 5 (9%) |
| Cirrhosis | | 4 (7%) |
| Diabetes mellitus | | 10 (18%) |
| Hypertension | | 23 (40%) |
| Prior myocardial infarction | | 4 (7%) |
| Congestive heart failure | | 5 (9%) |
| Peripheral vascular disease | | 1 (2%) |
| Chronic pulmonary disease | | 3 (5%) |
| Arthritis | | 4 (7%) |
| Peptic ulcer disease | | 2 (4%) |
| ***Medicines in last 24 hours* (2 variables)** | | |
| Use vasopressors/inotropes | | 34 (60%) |
| Use beta-blocker | | 6 (10%) |
| ***Alcohol & Smoking* (4 variables)** | | |
| Ever smoker | 7 (12%) | 33 (66%) |
| Current smoker | 7 (12%) | 20 (40%) |
| Cumulative smoking exposure (pack-years) | 9 (16%) | 20.96 ± 25.91 |
| AUDIT score | 7 (12%) | 6.58 ± 9.64 |
| ***Baseline vital signs* (8 variables)** | | |
| Heart rate (beats/min) | | 96.31 ± 18.78 |
| Systolic blood pressure (mmHg) | | 114.44 ± 23.34 |
| Diastolic blood pressure (mmHg) | | 58.30 ± 11.66 |
| Central venous pressure (mmHg) | 11 (19%) | 11.60 ± 5.24 |
| Mean arterial pressure (mmHg) | 7 (12%) | 73.20 ± 15.12 |
| Body temperature (°C) | | 37.31 ± 1.02 |
| Lowest respiratory rate (breaths/min) | | 17.74 ± 6.05 |
| Highest respiratory rate (breaths/min) | | 33.40 ± 11.41 |
| ***Baseline lab tests* (10 variables)** | | |

| | | |
|---|---|---|
| Hemoglobin (g/dL) | | 10.49 ± 2.27 |
| Serum sodium (mEq/L) | | 138.70 ± 4.80 |
| Serum potassium (mEq/L) | | 3.99 ± 0.65 |
| Log of serum glucose (mg/dL) | | 4.77 ± 0.34 |
| Serum bicarbonate (mEq/L) | | 24.26 ± 6.30 |
| Lowest hematocrit (%) | | 29.30 ± 6.23 |
| Log of lowest white blood cell count ($mm^3$) | | 8.91 ± 1.18 |
| Log of lowest platelet count (×1000/$mm^3$) | | 4.96 ± 0.77 |
| Log of highest blood urea nitrogen (mg/dL) | 1 (2%) | 2.99 ± 0.77 |
| Highest serum creatinine (mg/dL) | | 1.78 ± 1.48 |
| ***Baseline ventilator (8 variables)*** | | |
| Total respiratory rate (breaths/min) | | 23.65 ± 6.99 |
| Total minute ventilation (L/min) | | 10.86 ± 3.36 |
| Positive end-expiratory pressure (PEEP, cm $H_2O$) | | 8.58 ± 2.99 |
| Fraction of Inspired oxygen ($FiO_2$) | | 0.55 ± 0.19 |
| Oxygen saturation (SpO2, %) | | 96.32 ± 2.71 |
| Plateau pressure (cm $H_2O$) | 7 (12%) | 23.04 ± 5.78 |
| Peak inspiratory pressure (cm $H_2O$) | | 26.81 ± 7.54 |
| Mean airway pressure (cm $H_2O$) | 5 (9%) | 13.96 ± 4.31 |
| ***Primary outcome – Mortality*** **(3 variables)** | | |
| Days to death (days) | | 90.0 (90.0 - 90.0) |
| 60-day mortality | | 10 (18%) |
| 90-day mortality | | 12 (21%) |
| ***Secondary outcome – ICU index*** **(3 variables)** | | |
| Ventilator-free days to day 28 (days) | | 20.0 (6.0 - 23.0) |
| ICU-free days to day 28 (days) | | 18.0 (6.0 - 22.0) |
| APACHE III score | 1 (2%) | 87.41 ± 25.48 |
| ***Secondary outcome – Organ failure*** **(6 variables)** | | |
| Cardiovascular failure-free days to day 28 (days) | | 25.0 (20.0 - 26.0) |
| CNS failure-free days to day 28 (days) | | 19.0 (1.0 - 22.0) |
| Coagulation failure-free days to day 28 (days) | | 28.0 (26.0 - 28.0) |
| Renal failure-free days to day 28 (days) | | 28.0 (16.0 - 28.0) |
| Hepatic failure-free days to day 28 (days) | | 28.0 (25.0 - 28.0) |
| No. of days without failure of organs (days) | | 19.0 (7.0 - 25.0) |

Summary statistics were calculated on preprocessed clinical data, before the imputation of missing values. For continuous variables, statistics are reported as mean ± standard deviation. For categorical variables, statistics are reported as counts (percentage). Ventilator-free days and ICU-free days were recorded with deaths within 28 days assigned a value of zero, and statistics are reported as median (25th-75th percentile). Six organ failure indices were conditional on survival; days were counted only while patients were alive and were not assigned a value of 0 in cases of death within 28 days; Statistics are reported as median (25th-75th percentile). Missing counts and percentages are reported only when present; otherwise, the cells are left blank. Abbreviation: ARDS, acute respiratory distress syndrome; ICU, intensive care unit; CNS, central nervous system.

**Table S2**. Differentially expressed proteins identified by Wilcoxon rank-sum test

See file <**TableS2.csv**>

**Table S3**. Pneumotype-associated differentially expressed proteins identified by univariate logistic regression.

See file <**TableS3.xlsx**>

**Table S4**. Functional enrichment annotation groups obtained by hierarchical clustering of enriched GO and KEGG terms from pneumotype-associated differentially expressed proteins.

See file <**TableS4.xlsx**>

**Table S5**. Comparison of 90-day mortality and 28-day ICU variables between pneumotypes.

| Characteristic | Pneumotype 1 (n = 10) | Pneumotype 2 (n = 13) | Pneumotype 3 (n = 13) | Pneumotype 4 (n = 11) | Pneumotype 5 (n = 10) | *p*-value |
|---|---|---|---|---|---|---|
| 60-day mortality | 0 (0.00%) | 1 (7.69%) | 4 (30.77%) | 5 (45.45%) | 0 (0.00%) | 0.011 |
| 90-day mortality | 0 (0.00%) | 2 (15.38%) | 5 (38.46%) | 5 (45.45%) | 0 (0.00%) | 0.013 |
| Ventilator-free days | 20.0 (20.0 - 22.0) | 22.0 (10.0 - 23.0) | 14.0 (0.0 - 24.0) | 0.0 (0.0 - 24.0) | 21.0 (20.0 - 23.0) | 0.312 |
| ICU-free days | 19.5 (13.0 - 20.0) | 20.0 (7.0 - 22.0) | 11.0 (2.0 - 23.0) | 8.0 (0.0 - 23.0) | 19.5 (17.0 - 22.0) | 0.745 |
| APACHE III score | 77.49 ± 22.69 | 84.15 ± 22.76 | 101.85 ± 20.40 | 91.91 ± 32.55 | 77.50 ± 21.77 | 0.065 |

Summary statistics were calculated on clinical data with the imputation of the only one missing value in APACHE III score. Continuous variable APACHE III score is reported as mean ± standard deviation and compared using Welch's ANOVA. Ventilator-free days and ICU-free days are reported as median (25th-75th percentile) and compared using the Kruskal-Wallis *H* test. For 90-day mortality, statistics are reported as deaths (percentage) and compared using the Fisher's exact test due to expected cell counts in groups were < 5. Abbreviation: ARDS, acute respiratory distress syndrome; ICU, intensive care unit; ANOVA, analysis of variance.

**Table S6**. Comparison of organ failures within 28 day follow up between pneumotypes.

| Free days to day 28 (days) | Pneumotype 1 (n = 10) | Pneumotype 2 (n = 13) | Pneumotype 3 (n = 11) | Pneumotype 4 (n = 8) | Pneumotype 5 (n = 10) | *p*-value |
|---|---|---|---|---|---|---|
| Cardiovascular failure | 24.0 (23.0 - 25.0) | 25.0 (19.0 - 27.0) | 25.0 (21.0 - 26.0) | 24.0 (18.0 - 26.5) | 25.0 (21.0 - 28.0) | 0.880 |
| CNS failure | 22.0 (15.0 - 22.0) | 9.0 (1.0 - 22.0) | 6.0 (1.0 - 23.0) | 22.0 (11.5 - 22.0) | 15.5 (1.0 - 22.0) | 0.654 |
| Coagulation failure | 28.0 (27.0 - 28.0) | 28.0 (28.0 - 28.0) | 27.0 (22.0 - 28.0) | 28.0 (23.5 - 28.0) | 28.0 (28.0 - 28.0) | 0.127 |
| Renal failure | 28.0 (28.0 - 28.0) | 28.0 (27.0 - 28.0) | 26.0 (0.0 - 28.0) | 28.0 (21.0 - 28.0) | 28.0 (27.0 - 28.0) | 0.207 |
| Hepatic failure | 28.0 (27.0 - 28.0) | 28.0 (28.0 - 28.0) | 28.0 (25.0 - 28.0) | 28.0 (14.0 - 28.0) | 28.0 (27.0 - 28.0) | 0.792 |
| No organ failure | 24.0 (22.0 - 25.0) | 15.0 (7.0 - 24.0) | 11.0 (0.0 - 25.0) | 16.5 (4.5 - 26.5) | 21.5 (19.0 - 25.0) | 0.509 |

All organ injury indices were continuous but bounded (0 - 28); values are reported as median (25th-75th percentile) and compared using the Kruskal-Wallis *H* test. Five deaths (two in pneumotype 3 and three in pneumotype 4) were excluded from the table to account for death as a competing event. Abbreviation: ARDS, acute respiratory distress syndrome; CNS, central nervous system.

**Table S7**. Clinical baseline characteristics of the ARDS patients per pneumotype.

| Characteristic | Pneumotype 1 (n = 10) | Pneumotype 2 (n = 13) | Pneumotype 3 (n = 13) | Pneumotype 4 (n = 11) | Pneumotype 5 (n = 10) | *p*-value |
|---|---|---|---|---|---|---|
| ***Demographics* (5 variables)** | | | | | | |
| Age (years) | 45.20 ± 16.05 | 49.00 ± 14.15 | 54.92 ± 16.06 | 54.45 ± 20.71 | 41.40 ± 18.78 | 0.384 |
| Female | 3 (30%) | 4 (31%) | 7 (54%) | 7 (64%) | 3 (30%) | 0.347 |
| Race non-White | 0 (0%) | 2 (15%) | 4 (31%) | 4 (36%) | 4 (40%) | 0.162 |
| Ethnic Hispanic or Latino | 0 (0%) | 2 (15%) | 0 (0%) | 1 (9%) | 2 (20%) | 0.368 |
| Body mass index (kg/m²) | 26.53 ± 5.28 | 31.14 ± 7.17 | 27.19 ± 6.68 | 26.08 ± 5.99 | 24.39 ± 6.20 | 0.233 |
| ***ARDS severity* (2 variables)** | | | | | | |
| P/F ratio (mmHg) | 104.97 ± 25.39 | 113.75 ± 33.20 | 118.14 ± 47.73 | 157.13 ± 68.54 | 139.58 ± 59.01 | 0.173 |
| # of quadrants with opacities on chest X-ray | | | | | | 0.560* |
| 2 | 2 (20%) | 2 (15%) | 2 (15%) | 0 (0%) | 2 (20%) | |
| 3 | 2 (20%) | 2 (15%) | 1 (8%) | 4 (36%) | 4 (40%) | |
| 4 | 6 (60%) | 9 (69%) | 10 (77%) | 7 (64%) | 4 (40%) | |
| ***ARDS etiology* (6 variables)** | | | | | | |
| Trauma-related ARDS | | | | | | 0.385* |
| None | 10 (100%) | 11 (85%) | 12 (92%) | 11 (100%) | 8 (80%) | |
| Primary | 0 (0%) | 0 (0%) | 0 (0%) | 0 (0%) | 1 (10%) | |
| Secondary | 0 (0%) | 2 (15%) | 1 (8%) | 0 (0%) | 1 (10%) | |
| Sepsis-related ARDS | | | | | | 0.354* |
| None | 2 (20%) | 9 (69%) | 6 (46%) | 6 (55%) | 5 (50%) | |
| Primary | 4 (40%) | 2 (15%) | 3 (23%) | 1 (9%) | 2 (20%) | |
| Secondary | 4 (40%) | 2 (15%) | 4 (31%) | 4 (36%) | 3 (30%) | |
| Multiple blood transfusions | | | | | | 0.209* |
| None | 10 (100%) | 11 (85%) | 13 (100%) | 9 (82%) | 10 (100%) | |
| Primary | 0 (0%) | 1 (8%) | 0 (0%) | 1 (9%) | 0 (0%) | |
| Secondary | 0 (0%) | 1 (8%) | 0 (0%) | 1 (9%) | 0 (0%) | |
| Aspiration-related ARDS | | | | | | 0.336* |
| None | 7 (70%) | 10 (77%) | 10 (77%) | 7 (64%) | 4 (40%) | |
| Primary | 0 (0%) | 1 (8%) | 2 (15%) | 1 (9%) | 2 (20%) | |
| Secondary | 3 (30%) | 2 (15%) | 1 (8%) | 3 (27%) | 4 (40%) | |
| Pneumonia-related ARDS | | | | | | 0.519* |
| None | 1 (10%) | 4 (31%) | 6 (46%) | 6 (55%) | 4 (40%) | |
| Primary | 5 (50%) | 3 (23%) | 1 (8%) | 2 (18%) | 4 (40%) | |
| Secondary | 4 (40%) | 6 (46%) | 6 (46%) | 3 (27%) | 2 (20%) | |

| | | | | | | |
|---|---|---|---|---|---|---|
| Other lung injury | | | | | | 0.588* |
| None | 10 (100%) | 13 (100%) | 12 (92%) | 10 (91%) | 10 (100%) | |
| Primary | 0 (0%) | 0 (0%) | 0 (0%) | 1 (9%) | 0 (0%) | |
| Secondary | 0 (0%) | 0 (0%) | 1 (8%) | 0 (0%) | 0 (0%) | |
| ***Comorbidities* (14 variables)** | | | | | | |
| Baseline shock | 5 (50%) | 5 (38%) | 9 (69%) | 8 (73%) | 2 (20%) | 0.083 |
| Chronic dialysis | 0 (0%) | 1 (8%) | 1 (8%) | 1 (9%) | 1 (10%) | 0.999 |
| Non-Hodgkin's lymphoma | 0 (0%) | 1 (8%) | 0 (0%) | 0 (0%) | 0 (0%) | 0.999 |
| Metastatic solid tumor | 0 (0%) | 0 (0%) | 0 (0%) | 1 (9%) | 0 (0%) | 0.544 |
| Immunosuppression | 0 (0%) | 1 (8%) | 2 (15%) | 1 (9%) | 1 (10%) | 0.956 |
| Cirrhosis | 0 (0%) | 1 (8%) | 2 (15%) | 1 (9%) | 0 (0%) | 0.791 |
| Diabetes mellitus | 1 (10%) | 1 (8%) | 3 (23%) | 3 (27%) | 2 (20%) | 0.703 |
| Hypertension | 4 (40%) | 3 (23%) | 8 (62%) | 6 (55%) | 2 (20%) | 0.171 |
| Prior myocardial infarction | 0 (0%) | 1 (8%) | 1 (8%) | 2 (18%) | 0 (0%) | 0.632 |
| Congestive heart failure | 0 (0%) | 2 (15%) | 1 (8%) | 2 (18%) | 0 (0%) | 0.498 |
| Peripheral vascular disease | 0 (0%) | 0 (0%) | 0 (0%) | 1 (9%) | 0 (0%) | 0.544 |
| Chronic pulmonary disease | 0 (0%) | 1 (8%) | 1 (8%) | 1 (9%) | 0 (0%) | 0.999 |
| Arthritis | 1 (10%) | 1 (8%) | 1 (8%) | 1 (9%) | 0 (0%) | 0.999 |
| Peptic ulcer disease | 1 (10%) | 0 (0%) | 1 (8%) | 0 (0%) | 0 (0%) | 0.715 |
| ***Medicines in last 24 hours* (2 variables)** | | | | | | |
| Use vasopressors/inotropes | 5 (50%) | 5 (38%) | 10 (77%) | 9 (82%) | 5 (50%) | 0.137 |
| Use beta-blocker | 0 (0%) | 1 (8%) | 1 (8%) | 3 (27%) | 1 (10%) | 0.404 |
| ***Alcohol & Smoking* (4 variables)** | | | | | | |
| Ever smoker | 8 (80%) | 6 (46%) | 10 (77%) | 9 (82%) | 7 (70%) | 0.342 |
| Current smoker | 7 (70%) | 3 (23%) | 6 (46%) | 2 (18%) | 6 (60%) | 0.062 |
| Cumulative smoking exposure (pack-years) | 19.79 (6.00 - 20.38) | 0.00 (0.00 - 25.27) | 30.00 (10.00 - 45.00) | 20.00 (13.00 - 38.33) | 14.70 (0.00 - 30.24) | 0.273 |
| AUDIT score | 5.27 (3.00 - 12.00) | 0.00 (0.00 - 2.00) | 4.00 (0.00 - 8.00) | 1.00 (0.00 - 12.00) | 6.48 (0.00 - 14.00) | 0.114 |
| ***Baseline vital signs* (8 variables)** | | | | | | |
| Heart rate (beats/min) | 104.90 ± 14.34 | 93.08 ± 17.40 | 91.90 ± 22.29 | 91.55 ± 18.19 | 102.90 ± 18.82 | 0.242 |
| Systolic blood pressure (mmHg) | 103.90 ± 9.67 | 113.46 ± 25.04 | 109.00 ± 18.77 | 116.45 ± 22.69 | 131.10 ± 30.64 | 0.097 |
| Diastolic blood pressure (mmHg) | 53.50 ± 5.80 | 57.92 ± 11.38 | 55.85 ± 9.61 | 60.91 ± 15.15 | 63.90 ± 13.57 | 0.214 |
| Central venous pressure (mmHg) | 10.29 (10.00 - 12.00) | 12.00 (7.00 - 14.00) | 11.22 (9.00 - 15.00) | 11.61 (10.00 - 13.00) | 11.30 (9.00 - 12.79) | 0.941 |
| Mean arterial pressure (mmHg) | 71.50 (66.00 - 74.00) | 72.33 (68.00 - 78.00) | 74.00 (64.00 - 82.00) | 72.00 (67.00 - 81.00) | 78.04 (70.00 - 88.66) | 0.467 |

| | | | | | | |
|---|---|---|---|---|---|---|
| Body temperature (°C) | 37.96 ± 0.72 | 37.51 ± 0.81 | 36.98 ± 0.76 | 36.75 ± 1.66 | 37.43 ± 0.45 | 0.049 |
| Lowest respiratory rate (breaths/min) | 20.00 (15.00 - 22.00) | 18.00 (12.00 - 22.00) | 16.00 (13.00 - 18.00) | 22.00 (13.00 - 25.00) | 17.00 (15.00 - 18.00) | 0.360 |
| Highest respiratory rate (breaths/min) | 33.50 (30.00 - 46.00) | 32.00 (26.00 - 50.00) | 27.00 (20.00 - 31.00) | 32.00 (28.00 - 43.00) | 32.00 (24.00 - 40.00) | 0.206 |
| ***Baseline lab tests* (10 variables)** | | | | | | |
| Hemoglobin (g/dL) | 10.75 (10.20 - 11.40) | 9.80 (8.60 - 10.70) | 8.80 (8.60 - 10.60) | 10.40 (8.60 - 11.40) | 10.75 (9.50 - 12.30) | 0.452 |
| Serum sodium (mEq/L) | 136.80 ± 4.92 | 138.85 ± 3.51 | 140.15 ± 5.68 | 137.18 ± 4.17 | 140.20 ± 5.29 | 0.400 |
| Serum potassium (mEq/L) | 3.72 ± 0.21 | 3.95 ± 0.51 | 4.13 ± 0.95 | 4.19 ± 0.77 | 3.92 ± 0.43 | 0.166 |
| Log of serum glucose (mg/dL) | 4.73 (4.62 - 4.95) | 4.71 (4.61 - 4.84) | 4.92 (4.84 - 5.08) | 4.81 (4.61 - 5.26) | 4.70 (4.60 - 4.93) | 0.323 |
| Serum bicarbonate (mEq/L) | 25.00 (24.00 - 27.00) | 24.00 (19.00 - 31.00) | 24.00 (21.00 - 26.00) | 23.00 (15.00 - 25.00) | 24.00 (22.00 - 31.00) | 0.423 |
| Lowest hematocrit (%) | 32.70 ± 5.27 | 30.62 ± 7.79 | 26.62 ± 5.09 | 26.28 ± 5.53 | 31.00 ± 4.90 | 0.041 |
| Log of lowest white blood cell count ($mm^3$) | 9.44 (9.29 - 9.99) | 9.26 (8.96 - 9.53) | 8.79 (7.30 - 9.41) | 9.40 (8.79 - 9.71) | 9.07 (8.67 - 9.43) | 0.066 |
| Log of lowest platelet count (×1000/$mm^3$) | 4.83 ± 1.03 | 5.24 ± 0.53 | 4.83 ± 0.91 | 4.98 ± 0.72 | 4.86 ± 0.64 | 0.491 |
| Log of highest blood urea nitrogen (mg/dL) | 2.97 ± 0.71 | 2.83 ± 0.55 | 3.23 ± 0.93 | 2.86 ± 0.81 | 3.08 ± 0.85 | 0.734 |
| Highest serum creatinine (mg/dL) | 0.95 (0.70 - 1.40) | 1.10 (0.80 - 1.50) | 1.80 (1.10 - 2.50) | 1.20 (0.80 - 2.20) | 1.15 (0.90 - 4.10) | 0.371 |
| ***Baseline ventilator (8 variables)*** | | | | | | |
| Total respiratory rate (breaths/min) | 27.00 (22.00 - 30.00) | 22.00 (18.00 - 28.00) | 20.00 (18.00 - 24.00) | 30.00 (20.00 - 34.00) | 18.00 (17.00 - 30.00) | 0.168 |
| Total minute ventilation (L/min) | 12.33 ± 3.35 | 11.52 ± 3.74 | 9.30 ± 2.00 | 10.99 ± 4.28 | 10.42 ± 2.83 | 0.129 |
| Positive end-expiratory pressure (PEEP, cm $H_2O$) | 8.00 (5.00 - 10.00) | 8.00 (7.00 - 10.00) | 10.00 (8.00 - 12.00) | 10.00 (5.00 - 10.00) | 7.50 (5.00 - 10.00) | 0.282 |
| Fraction of inspired oxygen ($FiO_2$) | 0.53 (0.40 - 0.70) | 0.50 (0.40 - 0.70) | 0.50 (0.45 - 0.60) | 0.50 (0.40 - 0.60) | 0.50 (0.40 - 0.50) | 0.891 |
| Oxygen saturation (SpO2, %) | 97.00 (94.00 - 99.00) | 96.00 (95.00 - 98.00) | 97.00 (94.00 - 98.00) | 98.00 (95.00 - 100.00) | 96.00 (95.00 - 98.00) | 0.452 |
| Plateau pressure (cm $H_2O$) | 21.00 ± 5.12 | 23.78 ± 5.03 | 24.80 ± 6.62 | 23.26 ± 5.05 | 21.03 ± 5.09 | 0.438 |
| Peak inspiratory pressure (cm $H_2O$) | 25.50 ± 5.91 | 26.54 ± 7.87 | 29.46 ± 6.81 | 27.09 ± 10.01 | 24.70 ± 6.78 | 0.538 |
| Mean airway pressure (cm $H_2O$) | 12.90 ± 2.96 | 14.60 ± 5.08 | 15.58 ± 4.45 | 14.03 ± 4.11 | 12.30 ± 3.37 | 0.327 |

Summary statistics were calculated on preprocessed clinical data, with the imputation of missing values. Continuous variables with normal distribution across all pneumotypes (Shapiro-Wilk test $p > 0.05$ in each pneumotype) are reported as mean ± standard deviation and compared using Welch's ANOVA. Variables not normally distributed (Shapiro-Wilk test $p < 0.05$ in any pneumotype) are reported as median (25th-75th percentile) and compared using the Kruskal-Wallis *H* test. For nominal categorical variables, statistics are reported as counts (percentage) and compared using the Fisher's exact test due to expected cell counts in groups were < 5. $*$ Ordinal variables were analyzed using the Kruskal-Wallis *H* test instead of the Fisher's exact test. Abbreviation: ARDS, acute respiratory distress syndrome; ANOVA, analysis of variance.

**Table S8**. Comparison of alpha diversity of all detected microbial proteins between pneumotypes and controls.

| Alpha diversity | Pneumotype 1 (n = 10) | Pneumotype 2 (n = 13) | Pneumotype 3 (n = 13) | Pneumotype 4 (n = 11) | Pneumotype 5 (n = 10) | Controls (n = 49) | *p*-value |
|---|---|---|---|---|---|---|---|
| Species richness | 526 (468, 543) | 545 (542, 548) | 489 (465, 537) | 532 (479, 546) | 523 (473, 547) | 515 (456, 529) | <0.001 |
| Berger-Parker | 0.17 (0.14, 0.25) | 0.11 (0.10, 0.13) | 0.18 (0.13, 0.28) | 0.18 (0.13, 0.25) | 0.13 (0.09, 0.16) | 0.16 (0.12, 0.25) | 0.018 |
| Faith's phylogenetic diversity | 219,670 (202,230, 223,701) | 224,483 (222,780, 225,202) | 217,944 (209,462, 220,463) | 219,082 (212,118, 221,630) | 214,283 (212,368, 226,700) | 220,428 (210,219, 224,221) | 0.071 |
| Shannon | 4.43 (3.12, 4.45) | 4.52 (4.34, 4.57) | 4.15 (3.35, 4.32) | 4.20 (3.57, 4.50) | 4.35 (3.54, 4.42) | 3.80 (3.33, 4.14) | <0.001 |

Summary statistics were calculated on median taxonomic biomass proportions based on all detected microbial proteins. All continuous alpha diversity indices are reported as median (25th-75th percentile) and compared using the Kruskal-Wallis *H* test.

**Table S9**. Comparison of alpha diversity of 90 differentially expressed proteins across pneumotypes.

| Alpha diversity | Pneumotype 1 (n = 10) | Pneumotype 2 (n = 13) | Pneumotype 3 (n = 13) | Pneumotype 4 (n = 11) | Pneumotype 5 (n = 10) | *p*-value |
|---|---|---|---|---|---|---|
| Species richness | 59 (55, 61) | 63 (63, 65) | 59 (55, 63) | 60 (57, 62) | 61 (58, 65) | 0.003 |
| Berger-Parker | 0.25 (0.19, 0.29) | 0.20 (0.19, 0.24) | 0.32 (0.22, 0.53) | 0.35 (0.24, 0.37) | 0.30 (0.21, 0.39) | 0.019 |
| Faith's phylogenetic diversity | 74,735 (71,634, 76,536) | 79,924 (78,713, 81,629) | 77,666 (74,078, 80,352) | 77,985 (75,847, 81,782) | 79,087 (76,012, 80,967) | 0.031 |
| Shannon | 2.71 (2.60, 2.75) | 2.84 (2.74, 2.90) | 2.53 (2.14, 2.82) | 2.55 (2.18, 2.78) | 2.52 (2.29, 2.91) | 0.059 |

Summary statistics were calculated on median taxonomic biomass proportions based on 90 differentially expressed proteins. All continuous alpha diversity indices are reported as median (25th-75th percentile) and compared using the Kruskal-Wallis *H* test.

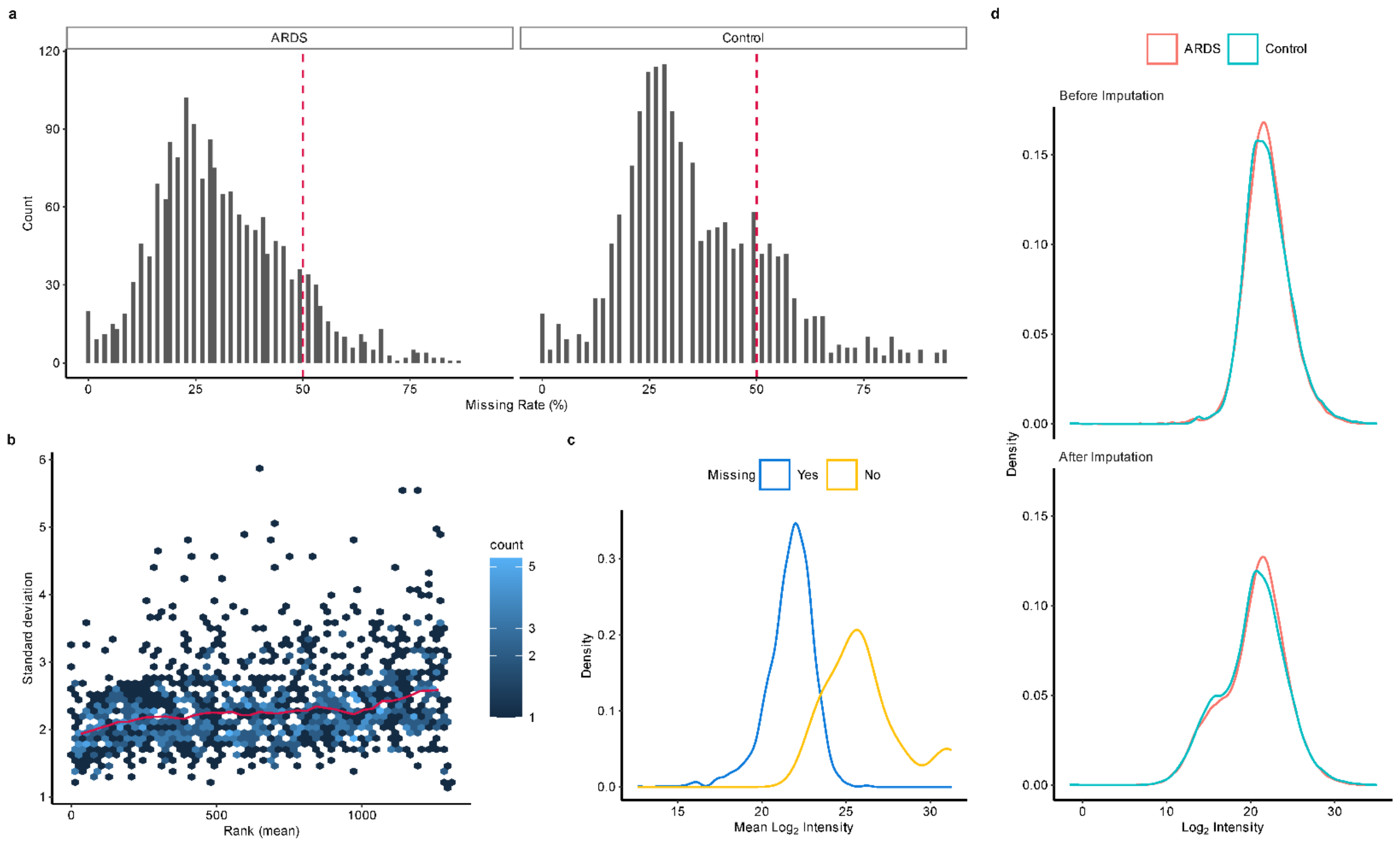


**Figure S1**. Filtering, normalization, and missing value imputation in microbial proteomic data. (a) Histogram of protein missing rates per sample in the ARDS and control groups, shown separately. The red dashed line indicates the 50% missingness threshold used for protein filtering. (b) Scatter plot of mean versus standard deviation after variance-stabilizing normalization. (c) Histogram of protein intensities in samples with versus without missing values, highlighting the systematic low-intensity pattern associated with missingness. (d) Histogram of protein intensities before and after left-censored imputation, shown separately for the ARDS and control groups. ARDS, acute respiratory distress syndrome.

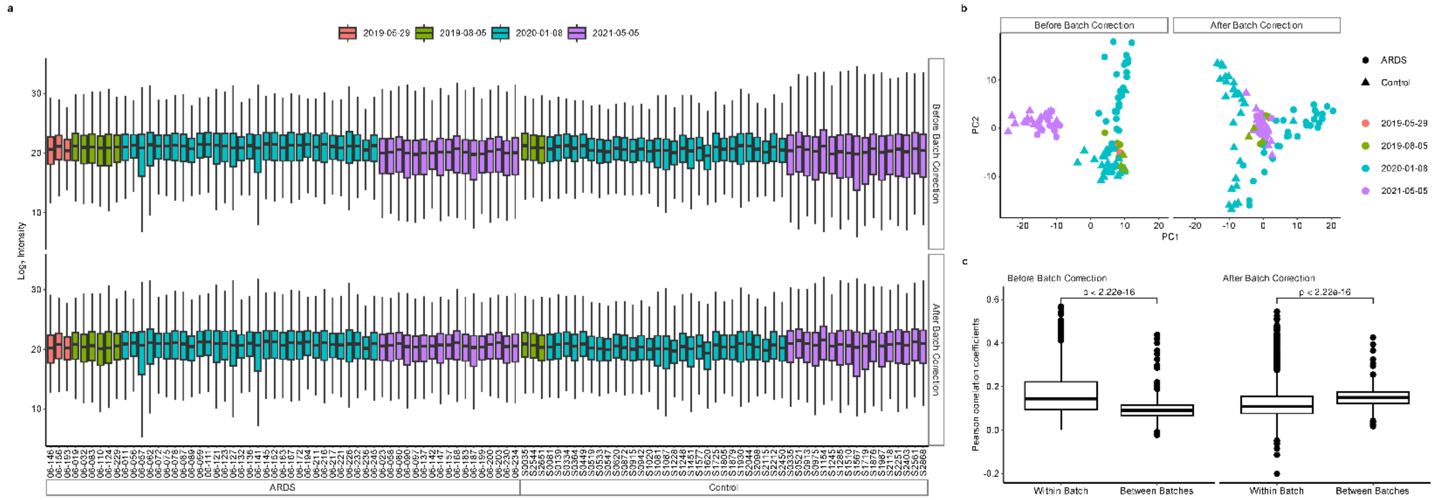


**Figure S2**. Batch effect adjustment in microbial proteomic data. (a) Boxplot of mean protein intensities per sample before and after batch effect correction, with samples colored by sequencing batch. (b) PCA plots of samples before and after ComBat-based batch adjustment, colored by sequencing batch, illustrating improved batch mixing after adjustment. (c) Boxplots of Pearson correlation coefficients for within-batch and between-batch sample pairs, shown before and after batch correction, demonstrating reduced batch-associated variability after adjustment. PCA, principal component analysis.

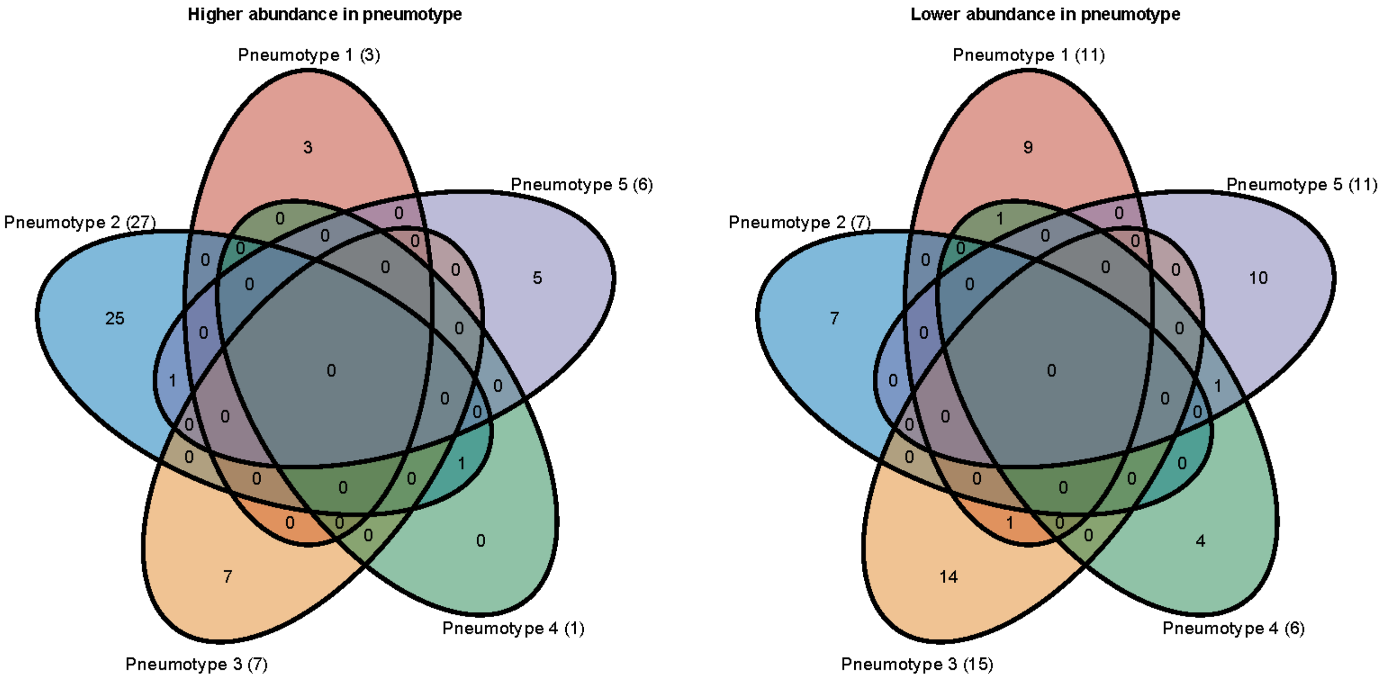


**Figure S3**. Venn diagrams of pneumotype-associated microbial differentially expressed proteins across five pneumotypes. For each pneumotype, pneumotype-associated proteins were identified using univariate logistic regression in a one-versus-rest framework without covariates; proteins with coefficient $p < 0.05$ were retained. The left panel shows pneumotype-associated proteins with positive coefficients (higher abundance in the pneumotype), and the right panel shows proteins with negative coefficients (lower abundance in the pneumotype).

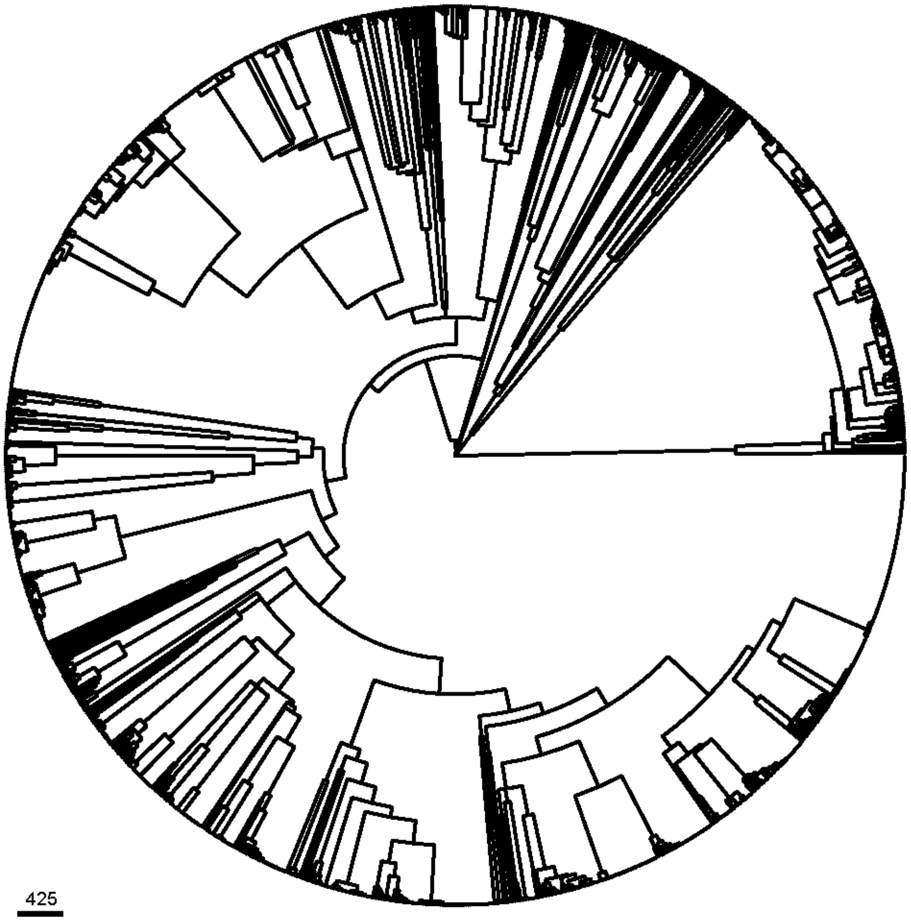


**Figure S4**. Phylogenetic tree constructed from species assigned to all detected microbial proteins in the metaproteomic dataset. Branch lengths correspond to estimated evolutionary divergence time, with the scale bar representing millions of years (Mya).

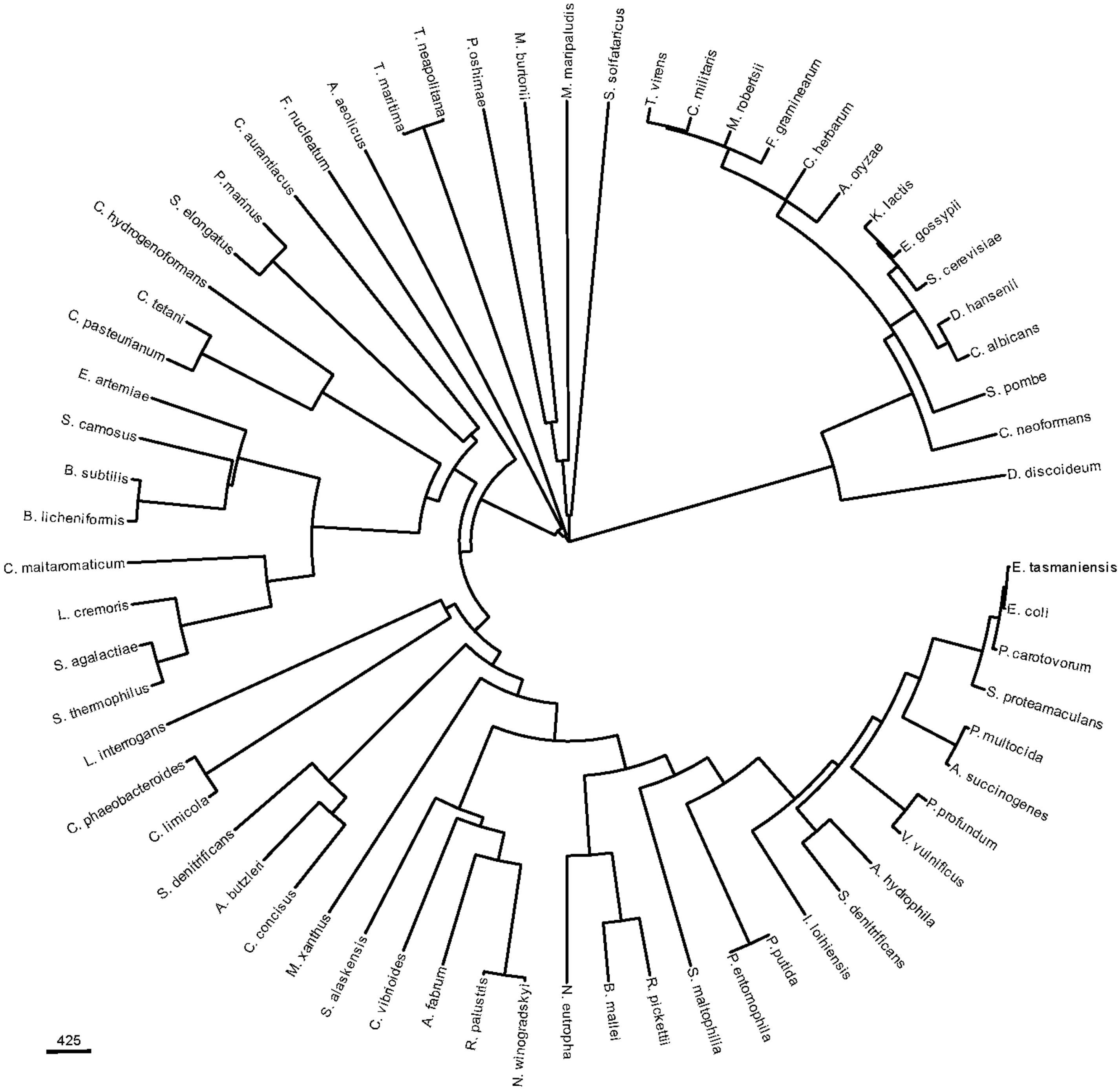


**Figure S5**. Phylogenetic tree constructed from species assigned to 90 differentially expressed proteins. Branch lengths correspond to estimated evolutionary divergence time, with the scale bar representing millions of years (Mya).

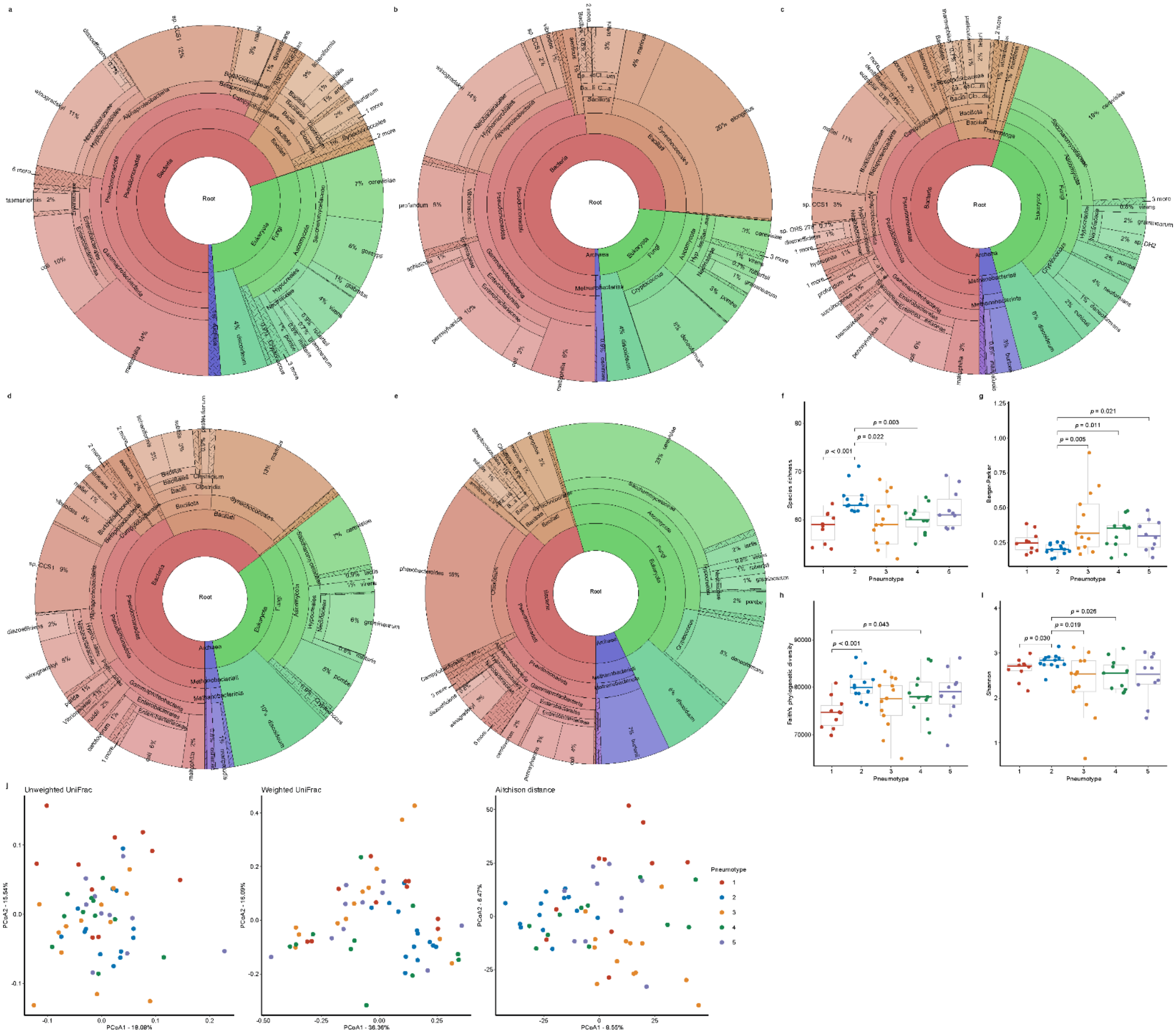

**Figure S6**. Characteristics of species assigned to differentially expressed proteins (DEPs) across pneumotypes. (a-e) Krona plots illustrating the taxonomic composition of species corresponding to DEPs across pneumotypes P1-P5. Proportions represent the relative taxon-attributed protein biomass contributions, calculated from summed protein intensities. Alpha diversity indices derived from the same species-level biomass proportions include (f) species richness, (g) Berger-Parker index, (h) Faith's phylogenetic diversity, and (i) Shannon diversity index. P-values were obtained using the Wilcoxon rank-sum test. (j) Beta diversity analysis, including two-dimensional principal coordinates analysis (PCoA) ordination based on unweighted (left) and weighted (middle) UniFrac distances (unique fraction metric) and Aitchison distance (right).
